# Parameter optimisation for mitigating somatosensory confounds during transcranial ultrasonic stimulation

**DOI:** 10.1101/2025.03.19.642045

**Authors:** Benjamin R. Kop, Linda de Jong, Kim Butts Pauly, Hanneke E.M. den Ouden, Lennart Verhagen

## Abstract

Transcranial ultrasonic stimulation (TUS) redefines what is possible with non-invasive neuromodulation by offering unparalleled spatial precision and flexible targeting capabilities. However, peripheral confounds pose a significant challenge to reliably implementing this technology. While auditory confounds during TUS have been studied extensively, the somatosensory confound has been overlooked thus far. It will become increasingly vital to quantify and manage this confound as the field shifts towards higher doses, more compact stimulation devices, and more frequent stimulation through the temple where co-stimulation is more pronounced. Here, we provide a systematic characterisation of somatosensory co-stimulation during TUS. We also identify the conditions under which this confound can be mitigated most effectively by mapping the confound-parameter space. Specifically, we investigate dose-response effects, pulse shaping characteristics, and transducer-specific parameters. We demonstrate that somatosensory confounds can be mitigated by avoiding near-field intensity peaks in the scalp, spreading energy across a greater area of the scalp, ramping the pulse envelope, and delivering equivalent doses via longer, lower-intensity pulses rather than shorter, higher-intensity pulses. Additionally, higher pulse repetition frequencies and fundamental frequencies reduce somatosensory effects. Through our systematic mapping of the parameter space, we also find preliminary evidence that particle displacement (strain) may be a primary biophysical driving force behind peripheral somatosensory co-stimulation. This study provides actionable strategies to minimise somatosensory confounds, which will support the thorough experimental control required to unlock the full potential of TUS for scientific research and clinical interventions.

**Highlights:**

- Tactile, thermal, and even painful somatosensory confounds may occur during TUS.
- Confounds can be mitigated via pulse shaping & transducer-specific parameters.
- Valid and replicable TUS research requires control for peripheral confounds.
- Particle displacement may be a primary driving force for somatosensory confounds.

## 1. Introduction

Transcranial ultrasonic stimulation (TUS) redefines the limits of non-invasive neuromodulation with its unprecedented spatial resolution and targeting capabilities^1–11^. However, peripheral co-stimulation poses a significant challenge to the reliable application of this technology. Peripheral effects such as somatosensation increase subject burden, can result in false inferences^12–16^, and contribute to the substantial placebo effects observed during brain stimulation^14,17–19^. Stringent experimental control is therefore required to infer direct neuromodulatory contributions to observed effects. While auditory confounds during TUS have been studied extensively^14,20–27^, possible somatosensory confounds remain unexplored. As TUS is increasingly applied at higher doses, with compact stimulation devices, and over the temples^28–33^, it will be critical to effectively manage somatosensory co-stimulation to ensure validity and specificity in this rapidly advancing field.

When an ultrasound beam is focused directly on the peripheral nervous system (PNS), such as the fingertip, tactile sensations can be felt. Here, ultrasound stimulates mechanoreceptors in the skin, including Merkel cells, Ruffini endings, Meissner corpuscles, and Pacinian corpuscles, which predominately innervate A-b fibres^34–38^. At higher doses, thermal and nociceptive sensations emerge, likely driven by the recruitment of higher threshold mechanoreceptors that innervate Type 1 A-δ and C fibres^34,37,39–42^. Peripheral somatosensation of ultrasound relies on mechanosensitive ion channels, including TRPV1, TRPA1, TREK-1, TRAAK, and Piezo channels. These channels not only play a critical role in the biological mechanism underlying TUS in the PNS, but are similarly implicated in TUS neuromodulation in the central nervous system (CNS)^39,43–50^.

The parameters of the ultrasound stimulation, such as the fundamental and pulsing frequencies, influence somatosensation. For instance, lower fundamental frequencies elicit stronger sensations^34–37,41,42,51–53^. Notably, certain parameters such as fundamental frequency differ in their relative strength of key biophysical effects like particle displacement and acoustic radiation force (ARF). Therefore, parameter mapping studies provide a valuable opportunity to elucidate the primary biophysical mechanisms underlying ultrasonic neuromodulation^34^. However, studies using peripherally focused ultrasound typically employ stimulation parameters distinct from those used during TUS, and do not fully explore the conditions relevant in the context of somatosensory co-stimulation during transcranial neuromodulation. Therefore, in the present study we investigate peripheral somatosensory effects across parameters relevant specifically to transcranial ultrasound for neuromodulation of the human brain.

In this pre-registered^54^ study, we bring TUS somatosensory confounds into focus by qualitatively characterising their nature and systematically mapping the confound-parameter space to identify avenues to minimise their impact. To ensure sufficient sensitivity to detect the effects of manipulating stimulation parameters, we intentionally operate under conditions that we expect will amplify somatosensory confounds. We further leverage this systematic investigation to explore the primary biophysical mechanisms of TUS that drive neuromodulation and provide preliminary evidence of particle displacement as a central biophysical mechanism. By putting forward actionable strategies to mitigate somatosensory confounds, we equip researchers with tools to optimise TUS studies for minimal burden and high inferential power, thus advancing TUS towards reliable and impactful applications across scientific, commercial, and clinical settings.

## 2. Methods

### 2.1. Participants

Twenty-nine participants were recruited, and twenty-five participants completed the study (14 female, 11 male, aged 25±4.3). Three participants were excluded after MRI intake, because the target sample size was achieved. One participant was excluded for psychological distress unrelated to TUS. All participants were free of psychiatric and neurological disorders, had no contraindications to brain stimulation, and provided informed consent. The study was approved by the Radboud University faculty of Social Sciences ethics committee (ECSW-2024-085) and conducted in accordance with the Declaration of Helsinki.

### 2.2. MRI

Both T1w and ultra-short echo time (UTE) MRI scans were acquired for each participant to support TUS neuronavigation and acoustic simulations^55,56^. See Supplementary Table 2 for sequence specifications.

### 2.3. Transcranial ultrasonic stimulation (TUS)

TUS was delivered using two NeuroFUS systems (supplier/support: BrainBox Ltd., Cardiff, UK; manufacturer: Sonic Concepts Inc., Bothell, WA, USA). Each of the four-channel radiofrequency amplifiers powered one piezoelectric transducer via an electrical impedance matching network. The experiment involved three transducers: a two-element 250 kHz transducer (250-2CH; serial number: CTX250-014, aperture diameter *d* = 45 mm, area = 15.90 cm^2^), a two-element 500 kHz transducer (500-2CH; serial number: CTX500-006, *d* = 45 mm, area = 15.90 cm^2^), and a four-element 250 kHz transducer (250-4CH; serial number: CTX250-026, *d* = 64 mm, area = 33.18 cm^2^; Fig. 1E). Detailed specifications for each transducer along with hydrophone measurements are reported in Supplementary Figs. 1-2 and Supplementary Table 1, in line with ITRUSST Standardised Reporting Guidelines^57^. Transducer performance was monitored across sessions^58,59^.

**Fig. 1.**
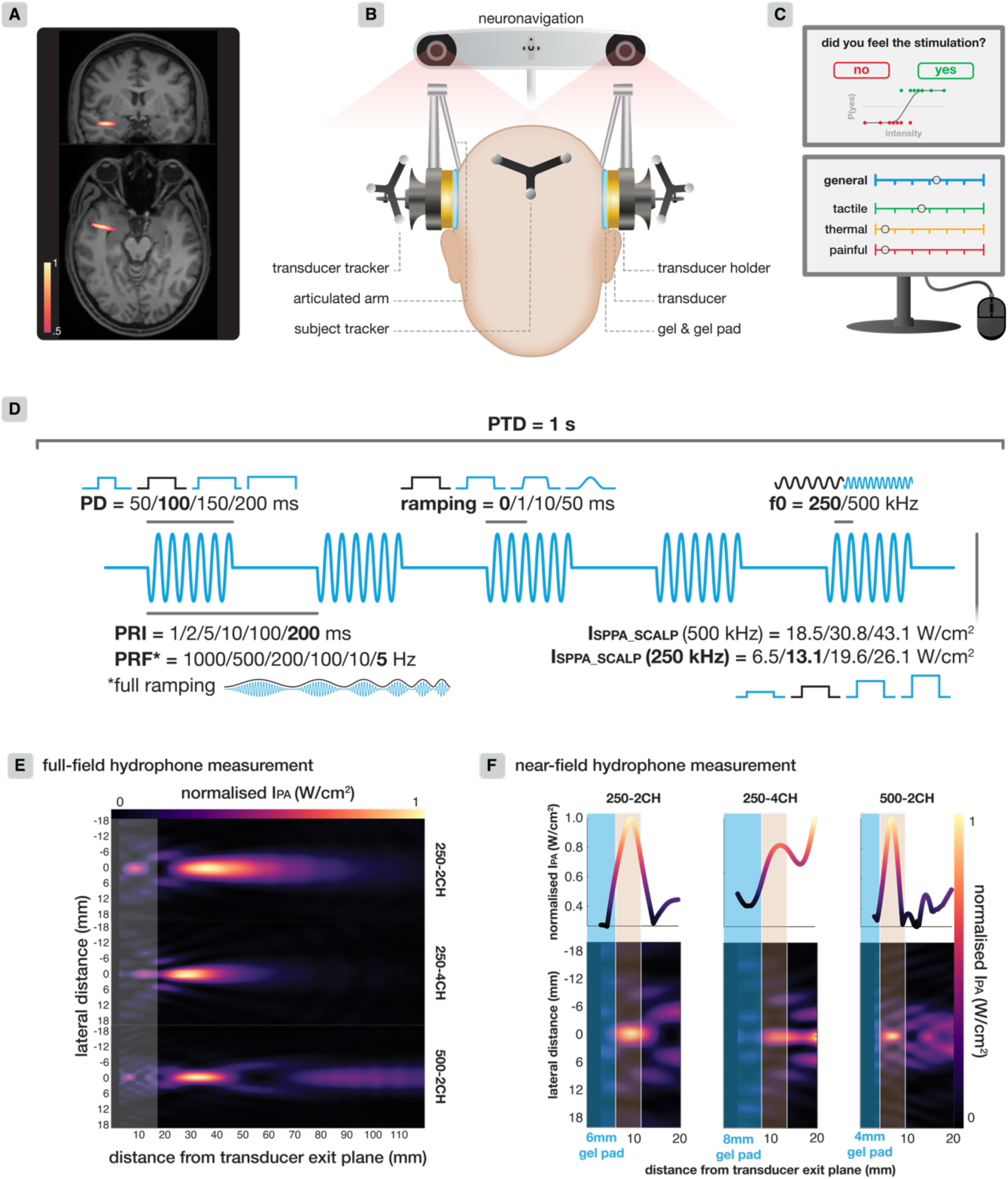
TUS experimental setup and methodology. **(A)** Representative acoustic simulation of temporal lobe white matter targeting, depicting the min-max normalised -3dB (full-width half-maximum) intensity profile for 250 kHz stimulation. **(B)** Experimental setup. **(C)** Experimental task. Top: yes/no questions used to estimate sensory thresholds (psychometric curve not visible to participant). Bottom: visual analogue scales. First, the overall holistic experience of somatosensory co-stimulation is captured with the ‘general’ VAS. Here, we determine whether TUS was felt only slightly, or very strongly. Next, subscales for tactile, thermal, and painful sensations capture the constituent sensory components specifically. **(D)** TUS protocol. Manipulated parameters are noted, with the standard protocol depicted in bold/black. Each parameter is manipulated separately while the other parameters remained standard with one exception: when investigating diUerent PRFs, full ramping was applied at each level. **(E)** Full-field hydrophone measurements for each transducer to quantify the intended transcranial acoustic field. The highlighted band reflects the transducer near-fields. **(F)** Near-field higher-resolution hydrophone measurements for each transducer, used to equalise exposure in the scalp. Gel-pad thickness (blue) for each transducer and the scalp (beige) are depicted. The gel-pad thicknesses, focal depths, and absolute stimulation intensities were adjusted such that the integrated maximum/total intensity in the scalp was equalised between transducers (see Supplementary Fig. 1-2 for details).

Transducers were coupled to the scalp using ultrasound gel^59^ and a gel-pad (Aquasonic 100 & Aquaflex, Parker Laboratories, NJ, USA). Prior to coupling, the participant’s hair around the stimulation site was prepared with ultrasound gel. Gel-pad thicknesses were 6, 8, and 4 mm for 250-2CH, 250-4CH, and 500-2CH transducers respectively, to optimise coherence of intensities in the scalp between transducers (Fig. 1F; see Supplementary Figs. 1-2 for details).

Transducer position was determined and maintained during the experiment by means of individualised neuronavigation based on participants’ T1w MRI scans (Localite GmbH, Sankt Augustin, Germany). TUS was targeted at the white matter of the temporal lobe – a region not expected to either produce or interact with sensory perception. A representative post-hoc acoustic simulation shows that temporal white matter targeting was successful (Fig. 1A; see Supplementary Fig. 3 for simulation methodology).

During the TUS experiment, two transducers were positioned bilaterally over the temporal window and held in place mechanically by articulated arms fastened to scaffolding built around the participant chair (Fig. 1B). A chin rest ensured that the participant was held firmly in place. Only one transducer administered stimulation per trial. To control for any putative difference in sensitivity to peripheral co-stimulation between the two sides of the head confounding observed differences between conditions, the transducer sides were switched halfway through the experiment, with the initial side counterbalanced between participants.

### 2.4. Somatosensory outcome measures

We quantified participants’ experience of TUS peripheral somatosensory co-stimulation through continuous visual analogue scales (VAS) and sensory thresholds. VAS ratings ranged from 0 (no sensation) to 10 (very intense sensation). First, participants provided an overall rating of their somatosensory experience of TUS (i.e., the ‘general’ rating scale). This initial rating captures the holistic perception of somatosensory co-stimulation intensity. Next, to gain more insight into the nature of the somatosensory effects, participants separately rated three subscales for tactile, thermal, and painful sensations specifically. This approach allowed us to capture both the overall experience of somatosensation, as well as to independently evaluate the nature of the sensations (Fig. 1C).

Sensory thresholds were measured by asking participants whether they could perceive a given protocol (yes/no) administered at various intensities. A custom thresholding procedure was used building on the parameter estimation by sequential testing method^60,61^ (see Supplementary Fig. 4 for details). The threshold was defined as the TUS intensity at which a fitted psychometric curve predicted a 50% likelihood of perception.

At the end of the experiment, we qualitatively characterised the somatosensory confound. Participants first responded to an open question prompting them to describe the sensations they experienced throughout the study. They then completed an adapted closed-format psychometric questionnaire for reporting somatosensory percepts^62^ (Fig. 2).

**Fig. 2.**
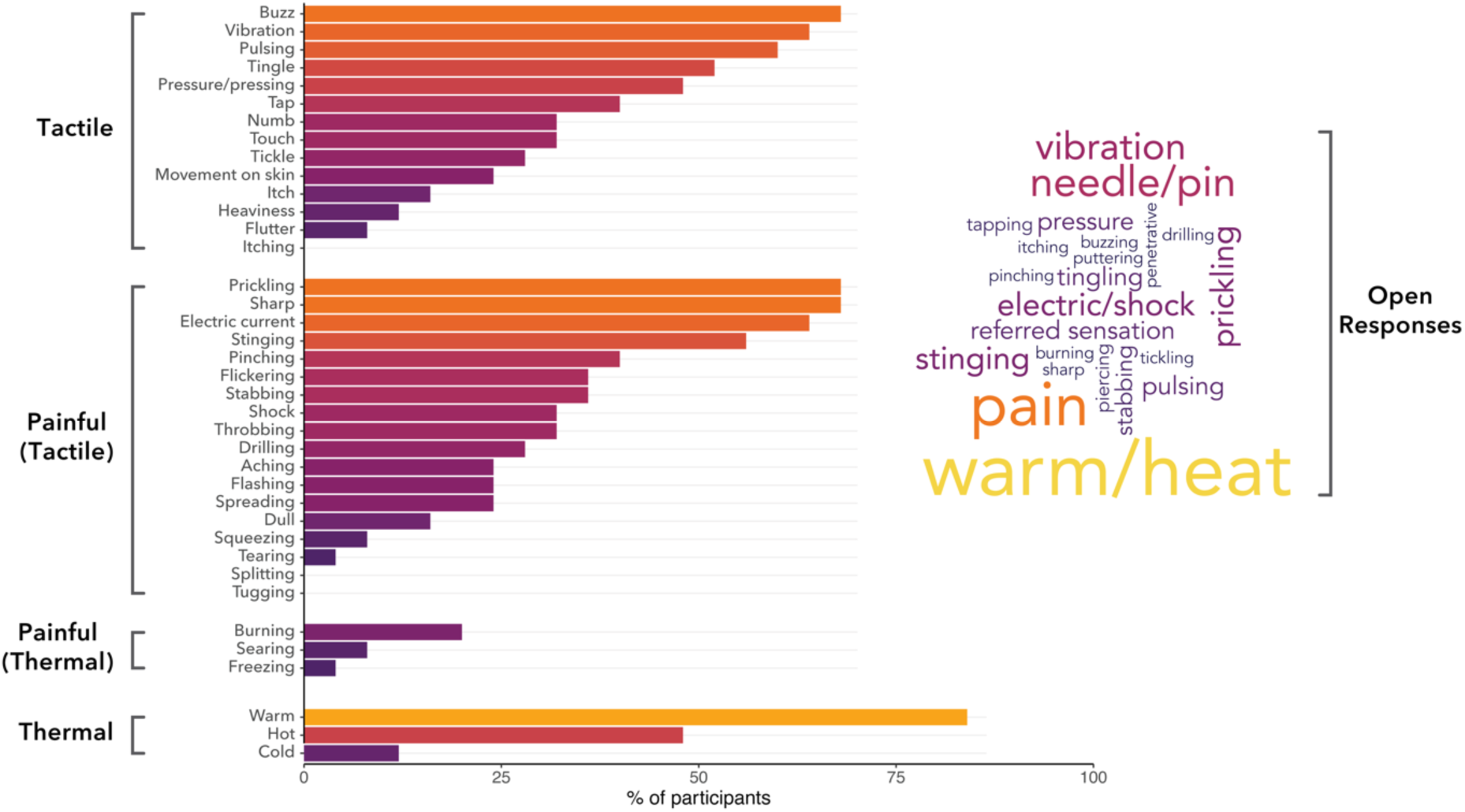
Characterisation of peripheral somatosensation during TUS. Descriptors were acquired via questionnaires completed after the main experiment. Here, participants retrospectively reported on all sensations they experienced across the entire session, encompassing all administered protocols collectively. Bars depict the percentage of participants who reported a given sensation on a closed psychometric questionnaire. The word cloud depicts descriptors mentioned in response to an open question, with size reflecting the frequency of a given descriptor.

**Fig. 3.**
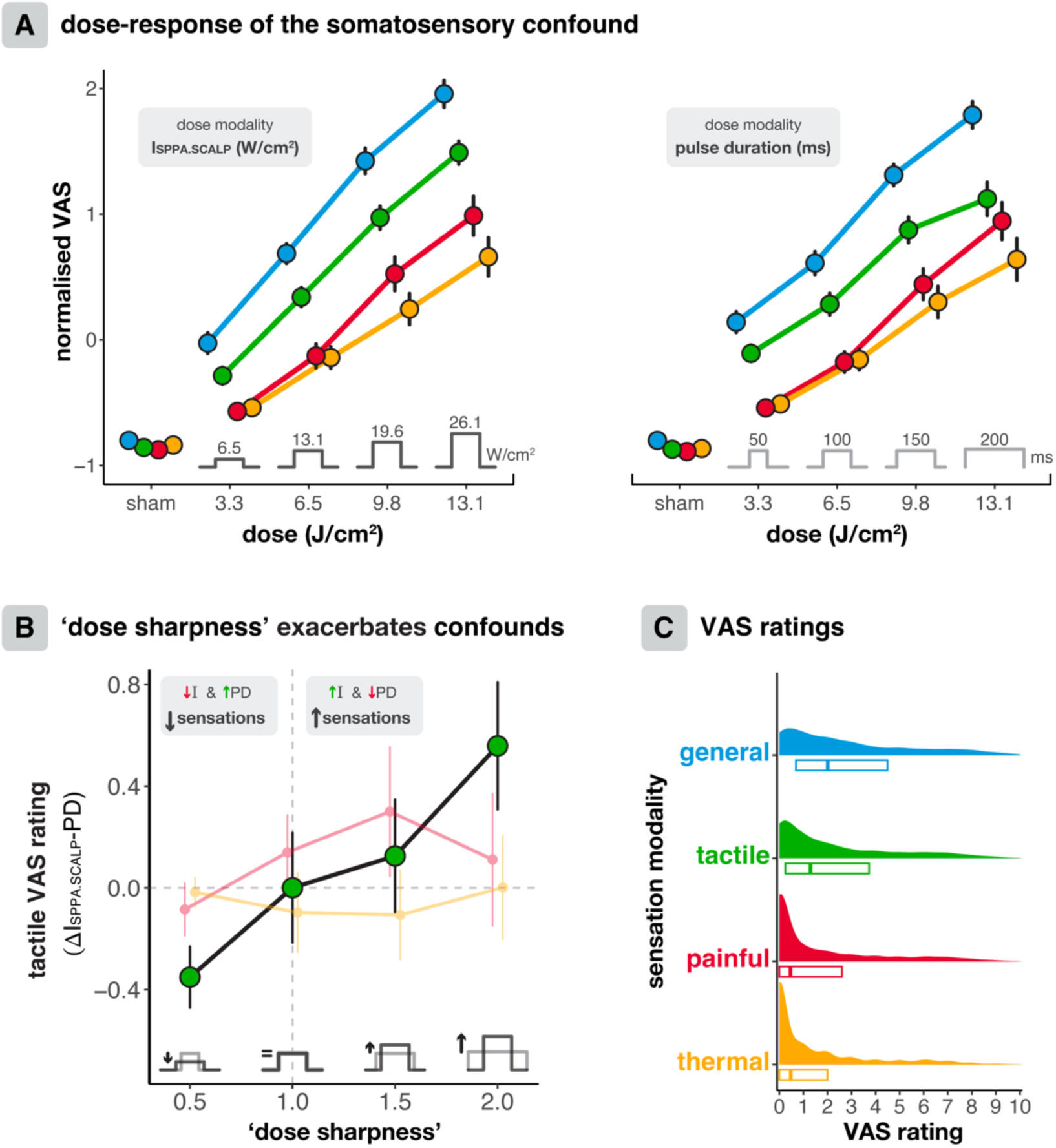
Dose-response of somatosensory confounds. **(A)** Dose-response of the somatosensory confound (250 kHz). Peripheral sensations become stronger as dose increases, both when increasing dose via intensity (left) and pulse duration (right) modalities. Tactile sensations are felt the earliest and the strongest. At higher doses, painful sensations become significantly stronger than thermal sensations. Points represent mean z-scored VAS ratings, error bars depict the standard error. **(B)** For tactile sensations specifically, higher ‘dose sharpness’ elicits stronger sensations. That is, shorter, higher intensity pulses cause more tactile sensations than longer, lower intensity pulses. Data reflect the absolute diMerence in VAS rating for the darker pulse waveform compared to the lighter pulse waveform. **(C)** Distribution of absolute VAS ratings across all conditions of the experiment, including participant ratings for the magnitude of co-stimulation they felt overall (i.e., general), as well as subscales for tactile, painful, and thermal sensations.

### 2.5. Study design

A sham-controlled, double-blind online TUS design was implemented, incorporating inter-subject trial-level counterbalancing. Full details on counterbalancing and task structure are provided in Supplementary Fig. 5.

The sham condition involved an auditory stimulus played over speakers, also present during TUS trials as an auditory mask. The volume was set uniformly across participants and was experienced as quite loud but not intolerable. This sound was designed to replicate the experiential qualities of our TUS protocols as closely as possible. Both the auditory stimulus and the code to generate it are publicly available here: https://doi.org/10.5281/zenodo.14052159. The PsychoPy^63^ IDE for Python was used to administer sham/auditory masking and TUS, set TUS parameters, and to record participants’ responses.

The standard TUS protocol (Fig. 1D, **bold**) was applied using transducer 250-2CH at a 250 kHz fundamental frequency (*f*_0_) with a square wave pulse repetition frequency (PRF) of 5 Hz, pulse repetition interval (PRI) of 200 ms, a pulse duration (PD) of 100 ms, a duty cycle (DC) of 50%, and a pulse train duration (PTD) of 1 second. The spatial-peak pulse-average intensity (I_SPPA_) was 19.72 W/cm^2^, and the corresponding near-field intensity in the scalp (I_SPPA.SCALP_) was 13.06 W/cm^2^. An inter-trial interval of approximately 10 seconds was used. All conditions adhered to ITRUSST recommendations for biophysical safety^64^ (see Supplementary Table 1 for safety metrics). To identify strategies to mitigate the somatosensory confound, we investigated multiple facets of this standard protocol, as well as transducer characteristics. Each section below describes a different investigation.

#### 2.5.1. Dose & dose modality (intensity/pulse duration)

We heuristically defined dose as exposure, that is, the integrated spatial-peak pulse-average intensity in the scalp (∫ *I_SPPA_SCALP_*) over the PTD. While recent discussions in the field distinguish between exposure and absorbed, equivalent, and effective dose^65^ – each accounting for interactions with biological tissue – we use the broader term ‘dose’ here for simplicity. Dose is given by the formula:

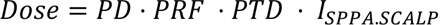

We applied stimulation at four doses: 3.3/6.5/9.8/13.1 J/cm^2^. The same dose was achieved via manipulation of two ‘dose modalities’: I_SPPA.SCALP_ and pulse duration (PD). We included dose modality to determine whether somatosensory co-stimulation was influenced by ‘dose sharpness’, i.e., the speed of equal dose delivery through shorter and higher intensity pulses versus longer and lower intensity pulses. Here, the interaction between ‘Dose’ and ‘Dose Modality’ (I_SPPA.SCALP_/PD) can yield insight into whether increasing intensity versus pulse duration has a different effect on the magnitude of somatosensory co-stimulation. Levels for I_SPPA.SCALP_ were 6.5/13.1/19.6/26.1 W/cm^2^, with PD = 100 ms held constant (‘PD_100ms_’). Levels for pulse duration were 50/100/150/200 ms, with I_SPPA.SCALP_ = 13.1 W/cm^2^ held constant (see Fig. 1D).

#### 2.5.2. Amplitude modulation (ramping)

We determined the effect of ramping on sensory thresholds by comparing square-wave modulated TUS with tapered cosine ramped amplitude modulation durations of 1 ms (0.01*PD_100ms_), 10 ms (0.1*PD_100ms_), and 50 ms (0.5*PD_100ms_; maximum ramping).

#### 2.5.3. PRF

PRFs were administered at 5/10/50/100/200/500/1000 Hz, covering the range of typically applied PRFs in the human literature to date^3,11,14,66^. Here, amplitude modulation consisted of a fully smoothed Tukey ramp (PD = PRI; tapered cosine ramp duration = 0.5⋅PRI), creating a more narrowband frequency distribution for the administered PRF, in contrast to the wider frequency distribution of square-wave pulse envelopes.

#### 2.5.4. Fundamental frequency (f0)

Stimulation was applied at 250 and 500 kHz using two transducers (250-2CH & 500-2CH). Focal depths, intensities, and gel-pad thicknesses were adjusted to optimise comparability and account for varying intensity (distribution) in the scalp between the two transducers (see Fig. 1F and Fig. 5). The dose-response relationship was mapped for 500 kHz, similar to 250 kHz, by manipulating the I_SPPA.SCALP_ (18.5/30.8/43.1 W/cm^2^; Supplementary Fig. 8).

#### 2.5.5. Transducer aperture area

The impact of aperture area on somatosensory co-stimulation was investigated by comparing two transducers with aperture areas of 15.90 and 33.18 cm^2^ (250-2CH & 250-4CH), each applying the same integrated total intensity to the scalp. The larger aperture transducer spread this energy over a wider area, thus reducing the intensity per unit area.

#### 2.5.6. Near-field peak amplitude

The annular arrays commonly used in human TUS research can produce near-field intensity peaks when the focus is steered axially. We quantified the impact of these near-field peaks on peripheral somatosensory co-stimulation by applying our standard TUS protocol at focal depth settings of 35.7, 38.3, 40.3 (standard), 42.1, and 44.1 mm. These depths corresponded to manufacturer-reported near-field peak intensities in the scalp of 5.3, 9.4, 13.8, 17.9, and 22.3 W/cm², respectively.

#### 2.5.7. Temporal summation

To examine whether somatosensory co-stimulation changes progressively throughout an online experiment, we tested three scenarios. First, we applied the standard protocol at approximately 1-minute intervals over a 10-minute segment of the experiment. Second, we applied the standard protocol in a series of six consecutive trials. Third, protocols were delivered in inter-subject counterbalanced sets, or “blocks”, allowing us to compare VAS ratings between consecutive sets of multiple TUS protocols (see Supplementary Fig. 5 for details).

To investigate possible sustained effects outlasting the stimulation period, as relevant for offline protocols with their longer pulse train durations (PTDs), we extended the PTD to 10 seconds at an I_SPPA.SCALP_ of 5.23 W/cm². Participants continuously reported their sensations on a VAS throughout this extended PTD, capturing the onset, development, and persistence of somatosensory co-stimulation in response to sustained stimulation.

### 2.6. Analysis

Data were processed, visualised, and analysed with R (v4.4.0). Data and code to reproduce the results will be provided following peer review. Sham-correction was performed by subtracting the average VAS rating for sham trials from each trial-level VAS rating per participant (see Supplementary Fig. 6 for sham). Linear mixed models (LMMs) were fitted to assess main effects and interactions for manipulated parameters on VAS ratings and sensory thresholds, typically with a maximal random effects structure^67^. These models were implemented through the lme4^68^ package in R. Statistical significance was set at a two-tailed α=0.05 and computed with t-tests using the Satterthwaite approximation of degrees of freedom.

For visualisation, VAS ratings were z-score normalised per participant to account for individual differences. These normalised data were used exclusively for visualisation to match the analyses they represent, as the linear mixed models we employed also account for this inter-individual variability.

## 3. Results

All participants reported feeling tactile, thermal, and painful sensations during the experiment. One participant experienced psychological distress unrelated to TUS and discontinued participation; their data was not analysed. Another participant displayed skin irritation at the stimulation site after participation, which resolved within a few hours. There were no further adverse events.

### 3.1. Qualitative characterisation of the somatosensory confound

On a closed psychometric questionnaire, more than half of participants reported warmth, buzzing, prickling, sharpness, electric current, vibration, pulsing, stinging, and tingling (Fig. 2). In response to an open question, the most prevalent sensations were warmth/heat, pain, a needle/pinprick, prickling, vibration, and electric current/shocks (Fig. 2). These sensations were measured after completion of the main experiment and therefore pertain to sensory experiences across the entire experiment. The co-occurrence of these sensations is depicted in Supplementary Fig. 7.

This somatosensory co-stimulation likely arises from direct stimulation of mechanoreceptors and sensory fibres, which are present in greater density at the temples as compared to the top of the scalp. TUS applied over the temples may additionally engage trigeminal ganglion cell bodies. Indeed, two participants reported referred sensations to their teeth and nose, respectively. All subsequent results we discuss pertain to the trial-by-trial VAS ratings (general/tactile/thermal/painful) during the main experiment.

Tactile sensations were rated significantly higher on the VAS than thermal and painful sensations for each applied intensity (Fig. 3; all *p* < 0.001). Thermal and painful sensations did not differ significantly for lower doses (i.e., 3.3 and 6.5 J/cm^2^), but painful sensations became significantly more salient than thermal sensations at higher doses (i.e., 9.8 and 13.1 J/cm^2^, all *p* < 0.001, Fig. 3A), potentially resulting from hyperactivation of receptor structures including those innervating A-b fibres^51,69^, or from central prioritisation of pain processing.

### 3.2. Dose-response relationship of somatosensory confounds

For both 250 and 500 kHz TUS, linear mixed models revealed that higher doses resulted in more peripheral somatosensation, as quantified by VAS ratings. At 250 kHz *f*_0_, four dose levels were tested (Fig 3B; 3/6/9/12 J/cm^2^). Dose was manipulated along two modalities, by increasing either intensity or pulse duration. A three-way LMM with a random intercept for Dose, Dose Modality, and Sensation Modality revealed a significant three-way interaction (*F*(2,4662) = 5.91, *p* = 0.003, η_p_^2^ = 0.003). Follow-up two-way LMMs with Dose and Dose Modality as predictors all revealed a significant main effect of Dose (all *p* < 0.0001). At a 500 kHz fundamental frequency (*f_0_*), there was a significant effect of Dose, manipulated solely though I_SPPA.SCALP_, for each Sensation Modality (Supplementary Fig. 8; all *p* < 0.001).

While these findings show that reducing dose can ameliorate the somatosensory confound across fundamental frequencies, it also poses a risk of diminishing the intended central nervous system neuromodulation. Importantly, our experiments also revealed opportunities to minimise the somatosensory confound while maintaining dose. For example, we found that ‘dose sharpness’, quantified as the ratio of peak intensity to duration at equivalent dose, predicts tactile somatosensory co-stimulation, which is the most prominent (Fig. 3B). Here, at equivalent doses, interactions reveal that shorter and higher intensity pulses cause more tactile somatosensation than longer and lower intensity pulses (Dose x Dose Modality: *F*(1,1522.2) = 15.5, *p* < 0.0001, η_p_^2^ = 0.01), with follow-up LMMs showing significant differences between Dose Modality for the lowest and highest conditions (Dose = 3.3: *F*(1,374) = 7.03, *p* = 0.008, η_p_^2^ = 0.018; Dose = 13.1: *F*(1,373) = 10.3, *p* = 0.001, η_p_^2^ = 0.027). We note that this relationship was significant for tactile sensations but was absent for painful and thermal sensations. Here, dose sharpness is experienced as ‘tapping’ rather than pain or heat. Thus, tactile sensations can be minimised while maintaining dose by favouring longer pulses with lower intensities over short pulses with higher intensities.

### 3.3. Pulse shaping & temporal characteristics

#### 3.3.1. Amplitude modulation (ramping)

Ramping significantly decreased the somatosensory confound (Fig. 4A; *F*(3,72) = 8.46, *p* < 0.0001, η_p_^2^ = 0.261), resulting in less sensation in response to 10 and 50 ms of tapered cosine amplitude modulation compared to square wave modulation (10 ms: *p* = 0.023; 50 ms: *p* < 0.0001; FDR corrected for multiple comparisons). In line with our findings for ‘dose sharpness’, this result suggests that the gradient of change in TUS amplitude may contribute to tactile peripheral co-stimulation.

**Fig. 4.**
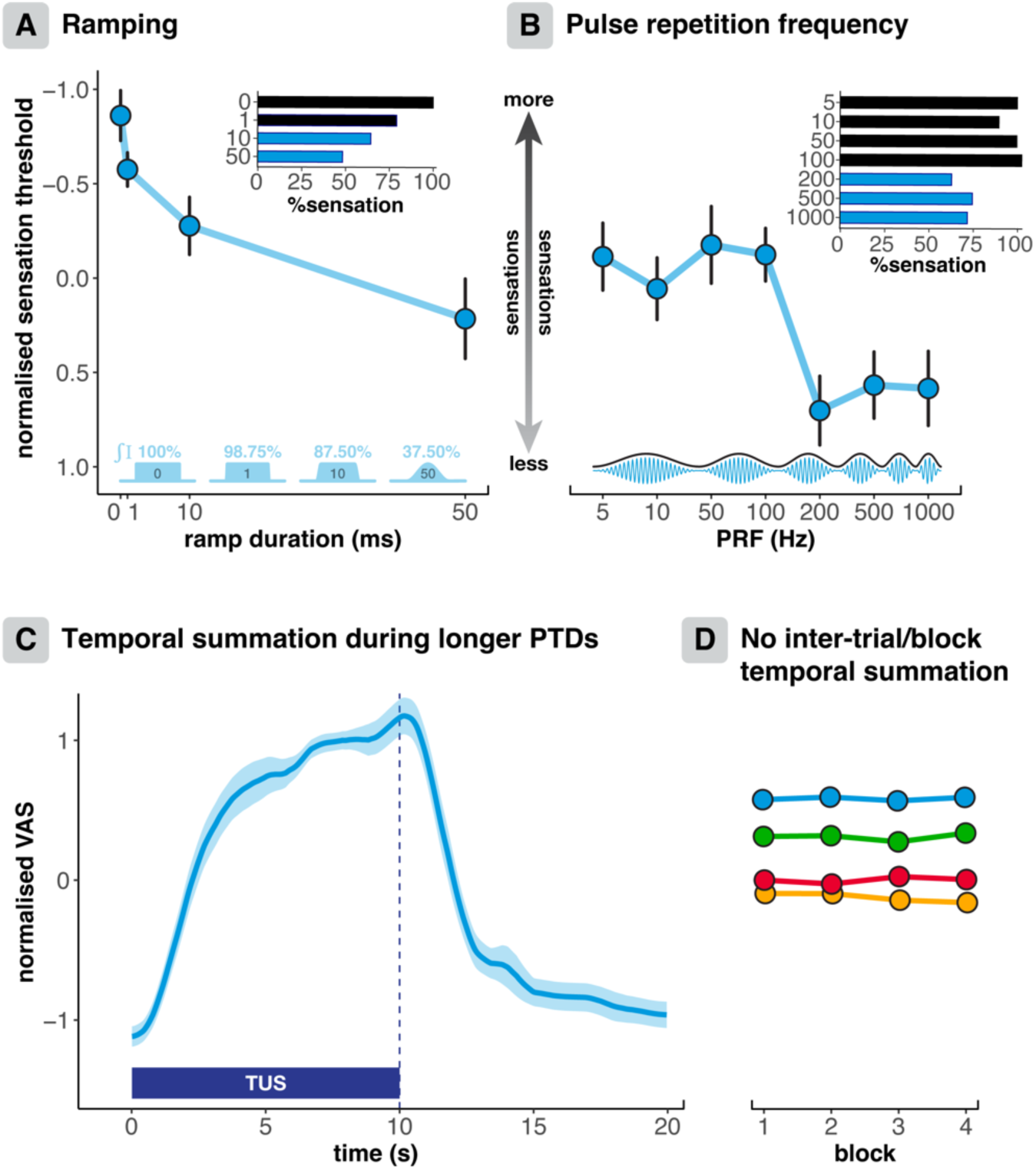
Pulse shaping & temporal characteristics. **(A)** Ramping for 10 and 50 ms significantly reduced the somatosensory confound. Normalised thresholds are depicted on a flipped y-axis, where visually higher points reflect stronger sensations (i.e., lower thresholds). The histogram depicts the average threshold as a percentage, and the pulse envelopes illustrate the integrated intensity as a percentage, both as compared to a square-wave pulse. **(B)** Higher pulse repetition frequencies (PRFs) elicited significantly less somatosensory co-stimulation than lower PRFs. **(C)** After a sharp initial incline, somatosensory co-stimulation increases steadily during a 10 second PTD, indicating that somatosensory confounds may develop over the longer PTDs typically used in oUline protocols. **(D)** Somatosensory co-stimulation remains constant across repeated sets (‘blocks’) of protocols, demonstrating that there is no inter-trial temporal summation of somatosensory co-stimulation in this online protocol. Additionally, there was no inter-trial temporal summation for identical protocols applied consecutively, nor when interspersed throughout the experiment (see Supplementary Fig. 9). Points represent condition means and error bars depict the standard error.

**Fig. 5.**
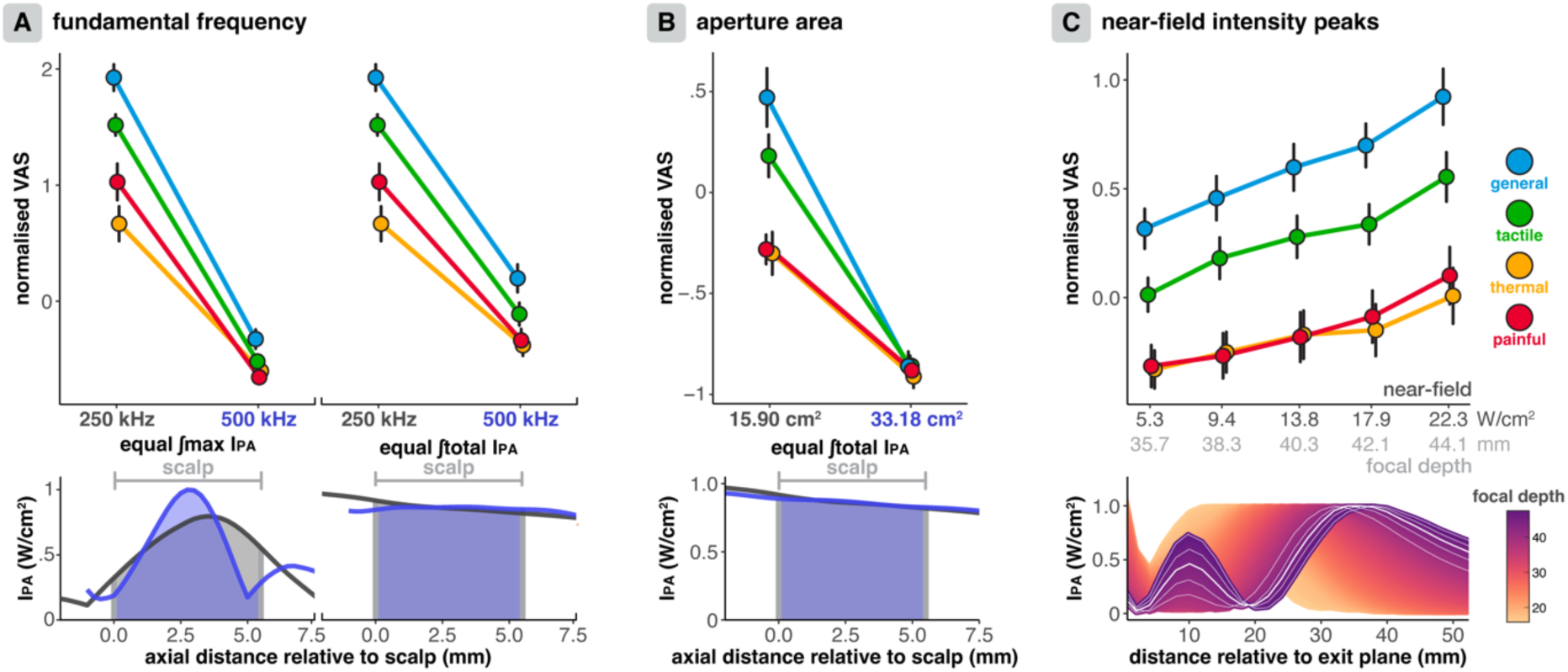
Transducer-specific parameters. **(A)** Higher fundamental frequencies elicit significantly less somatosensory co-stimulation, both when the integrated maximum intensity (bottom left) and integrated total intensity (bottom right) in the scalp are equalised. **(B)** A larger aperture area transducer delivering an equal integrated total intensity (bottom) also elicited less co-stimulation. **(C)** Higher magnitude near-field intensity peaks caused by axial steering to larger focal depths for this transducer elicited more somatosensory co-stimulation. On the lower panel, white lines indicate the manufacturer axial profile measurements for the applied focal depths. Points indicate conditions means and error bars depict standard error.

#### 3.3.2. Pulse repetition frequency (PRF)

Pulse repetition frequencies of 100 Hz and lower were associated with more peripheral somatosensation (i.e., lower thresholds) than higher pulse repetition frequencies (Fig. 4B). There was a significant main effect of PRF on thresholds (*F*(6,144) = 4.10, *p* = 0.001, η_p_^2^ = 0.146; see Supplementary Table 3 for post-hoc paired comparisons). Sensations for grouped PRFs of 5, 10, 50, and 100 Hz were significantly lower than for PRFs of 200, 500, and 1000 Hz (*F*(1,24) = 13.6, *p* = 0.001, η_p_^2^ = 0.361). These results suggest that peripheral sensory nerves are preferentially activated by neurophysiologically relevant PRFs within their endogenous firing rates.

#### 3.3.3. Temporally summative somatosensory co-stimulation

##### 3.3.3.1. No inter-trial cumulation of somatosensory confounds in an online paradigm

VAS ratings remained stable throughout this online experiment for a single stimulation protocol interspersed throughout a 10-minute stimulation period (Trial: *F*(1,24) = 0.061, *p* = 0.808, η_p_^2^ = 0.003, BF01 = 30.8), and for six successive trials of the same protocol (*F*(1,24) = 0.427, *p* = 0.52, η_p_^2^ = 0.017, BF01 = 28.1; see Supplementary Fig. 9). These results also demonstrate the consistency of the VAS ratings as an outcome measure. When assessing blocks of trials in which the same set of protocols were administered, VAS ratings also remained consistent over time (*F*(1,24) = 1.63, *p* = 0.214, η_p_^2^ = 0.064, BF01 = 283.6). In this specific but representative online experimental paradigm (ITI = 10 s, PTD = 1 s), there was no inter-trial cumulation of the somatosensory confound.

##### 3.3.3.2. Cumulation of somatosensory co-stimulation for o:line paradigms

Offline TUS protocols are typically characterised by longer pulse train durations where somatosensory co-stimulation may develop as stimulation progresses. Here, we show that following participants’ initial response to peripheral co-stimulation, sensations continue to build steadily until the stimulation ends, whereafter sensations subside almost immediately. Some minor sensory effects persist for a few seconds before returning fully to baseline.

### 3.4. Transducer-specific characteristics

#### 3.4.1. Fundamental frequency & biophysical mechanisms

TUS at a 500 kHz *f*_0_ elicited significantly less somatosensory co-stimulation than a 250 kHz *f0*, regardless whether the maximum or total integrated intensity in the scalp was equalised (Fig. 5A; ∫max.: *F*(1,24) = 97.6, *p* <0.0001, η_p_^2^ = 0.803; ∫total: *F*(1,24) = 58.7, *p* <0.0001, η_p_^2^ = 0.71). Therefore, increasing *f*_0_ can decrease somatosensory confounds.

We note that the direction of this relationship suggests that particle displacement may be a primary biophysical mechanism driving peripheral neuromodulation, as this biophysical effect is stronger at lower fundamental frequencies^70^. To further test this hypothesis, we investigated whether somatosensory co-stimulation scaled with pressure (∼particle displacement) or intensity (∼ARF). Specifically, we compared non-nested LMMs using intensity (*I*) or pressure 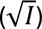 as a predictor. Across all Sensory Modalities, we found that pressure was >90% likely to better explain the variance in our data than intensity (general: ΔAIC = 10.8, *w* = 0.995; tactile: ΔAIC = 9.7, *w* = 0.992; thermal: ΔAIC = 5.9, *w* = 0.949; painful: ΔAIC = 4.9, *w* = 0.921).

#### 3.4.2. Transducer aperture area

A larger aperture area four-element annular array transducer (33.18 cm^2^) elicited significantly less somatosensory co-stimulation than a smaller two-element transducer (15.90 cm^2^) when equalising the integrated total intensity in the scalp (Fig 5B; *F*(1,24) = 40.5, *p* < 0.0001, η_p_^2^ = 0.628). Decreasing the intensity per unit area in the near-field using larger aperture transducers can maintain transcranial intensities while minimising peripheral somatosensory confounds.

#### 3.4.3. Near-field peaks

The amplitude of near-field peaks in the scalp – sometimes caused by axial steering with commonly used annular arrays – significantly impacts peripheral somatosensory co-stimulation (*F*(1,24) = 37.9, *p* < 0.0001, ηp^2^ = 0.612). Greater near-field peak amplitudes present at higher focal depths for this transducer (250-2CH) resulted in more somatosensation (Fig. 5C). These sensations could be eliminated by minimising near-field peaks in the scalp through improved transducer manufacturing and/or using an appropriate combination of axial steering and coupling medium offset.

## 4. Discussion

In this pre-registered^54^ study, we present evidence of peripheral somatosensory confounds during TUS in humans. A comprehensive understanding of the nature of these confounds and the conditions under which they arise is necessary to conduct well-controlled, robust, and minimally burdensome TUS research. Therefore, we systematically mapped the confound parameter space and demonstrated that somatosensory co-stimulation can be minimised by avoiding near-field peaks in the scalp, spreading energy across a greater area of the scalp, using ramped pulses, lowering ‘dose-sharpness’, and administering higher pulse repetition frequencies, higher fundamental frequencies, and lower doses. We also identify particle displacement as a putative biophysical driving force behind peripheral somatosensory confounds. With appropriate mitigation strategies, somatosensory co-stimulation can be minimised while maintaining meaningful TUS doses. Our findings lay the foundation for TUS parameter optimisation to enhance specificity and reliability in research and clinical settings.

### 4.1. Dose-response of the somatosensory confound

All participants experienced tactile, thermal, and painful sensations, with common descriptors including ‘buzzing’, ‘prickling’, ‘sharpness’, and ‘electric current’ (Fig. 2). Note that any noxious sensations are not caused by biological damage, and these sensations not present for all protocols. The primary determinant of somatosensory confounds is dose, defined as the integral of intensity over the pulse train^70^. This definition also aligns with the term ‘exposure’, whereas a more precise account of dose (e.g., absorbed, equivalent, or effective) would consider interactions with biological tissues (see Nandi et al., 2025)^65^. Nonetheless, the broader term ‘dose’ is used here for simplicity. Higher doses amplify the somatosensory confound, both when the achieved by increasing intensity or pulse duration, suggesting that these modalities are, to some extent, interchangeable components of dose.

Without adequate controls, observed effects may be erroneously attributed to transcranial neuromodulation, while their causative origin lies in peripheral confounds. Indeed, such misinterpretations have already arisen in the context of the TUS auditory confound^14^. The challenge, then, lies in determining how best to minimise this confound without simply reducing dose, which risks compromising the intended neuromodulatory effects in the brain. Multiple strategies for addressing this challenge are discussed below.

### 4.2. Pulse shaping & temporal characteristics

Reducing ‘dose sharpness’ by delivering an equivalent dose with longer, lower-intensity pulses instead of shorter, higher-intensity pulses effectively minimises tactile co-stimulation, which is the most prominent somatosensory confound (Fig. 3). Note that there likely remains an absolute minimum intensity required for neuromodulation, and excessively high duty cycles may negate certain pulse repetition frequency (PRF) related effects^71,72^. Nonetheless, several studies demonstrate that increasing dose through longer pulse durations can also reliably produce robust TUS effects^27,71–73^, and this therefore constitutes one avenue for somatosensory confound mitigation.

In addition, ramping the pulse envelope can effectively reduce somatosensory confounds by more than 50% (Fig. 4A). Specifically, we show that 10 and 50 ms tapered cosine ramps significantly increase sensory thresholds. However, it is important to note that ramping inherently reduces dose by lowering the intensity integral in proportion to the ramp length. There may be a trade-oa wherein the benefits of confound minimisation become outweighed by the reduction in dose beyond a given tipping point. In the present study, a 10 ms ramp duration offered the optimal balance: confounds were reduced by ∼45%, while dose was only marginally reduced by 12.5%, compared to a square-wave envelope. Previous studies have demonstrated that effective CNS neuromodulation remains feasible with ramped pulses^8,27,74–76^, thus supporting the viability of ramping for mitigation of somatosensory confounds, in addition to its well-established efficacy for auditory confounds^24,26,27,77^.

Pulse repetition frequencies (PRFs) of 200 Hz and higher elicited ∼30% less sensations than lower frequencies, suggesting that PRF can be tuned to minimise the somatosensory confound (Fig. 4B). Importantly, the dependence of co-stimulation magnitude on PRF suggests a relationship to endogenous neurophysiological firing rates^74,78^. For example, perhaps Type 1 rapidly adapting mechanoreceptors were preferentially activated in this study, given their sensitivity to the lower range of the applied PRFs^79,80^. Different PRFs may elicit distinct effects in relation to mechanoreceptive frequency sensitivity and firing rates^36^. Although higher PRFs can reduce co-stimulation, they are associated with stronger auditory confounds and offer limited opportunities for ramping, which our results suggest could be a more effective mitigation approach. Nonetheless, PRF can be considered as one of several parameters that can be optimised, with higher PRFs remaining capable of eliciting convincing neuromodulatory effects^11,81^.

Over longer timescales, somatosensory co-stimulation may develop progressively. Indeed, we show that longer pulse train durations, commonly used in offline TUS protocols, can elicit a gradual buildup of co-stimulation. Dividing these protocols into segments could mitigate this effect. For example, intermittent TUS protocols, such as the ‘accelerated theta-burst’ protocol, successfully incorporate 30-minute intervals between pulse trains^82^.

In contrast, there was no inter-trial cumulation of co-stimulation throughout this online experiment. Bayesian analyses strongly indicated equivalence in VAS ratings for identical protocols delivered successively or interspersed throughout trials, and between consecutive sets of protocols. The absence of inter-trial cumulation validates the feasibility of trial-based study designs where conditions are repeated over time.

### 4.3. Transducer-specific parameters

Intensity peaks in the transducer near-field can significantly contribute to peripheral co-stimulation and should be circumvented. These near-field peaks are common for the transducers widely employed in human TUS research, such as the annular arrays used in this study. Our findings show that, as the focal depth for our annular array transducer increased, so did both near-field peaks and somatosensory confounds (Fig 5C, lower panel). For currently available transducer designs, it is crucial to consider the transducer-specific focal depths at which these peaks occur and ensure they do not overlap with peripheral nerve structures. This can be achieved by selecting an appropriate combination of focal depth and coupling medium thickness.

The spread of intensity across the scalp can also be exploited to minimise confounds. Specifically, we show that a 33.18 cm^2^ aperture area transducer evoked substantially less somatosensation than a 15.9 cm^2^ aperture area transducer delivering the same integrated total intensity in the scalp. Multi-transducer constellations and hemispheric arrays^7–9,83,84^ can therefore also be expected to circumvent peripheral somatosensory confounds, ostensibly up to very high transcranial intensities, without a substantial impact on CNS neuromodulation.

Finally, higher fundamental frequencies (500 kHz) produced fewer somatosensory confounds than lower frequencies (250 kHz), even when integrated maximum and total intensities in the scalp were equalised. Importantly, the intensity in the brain was higher for 500 kHz stimulation, thus demonstrating that higher frequencies maintain their advantage in reducing somatosensory confounds even if considering differences in acoustic transmission. Reduced co-stimulation compared to 250 kHz could be influenced by factors including smaller near-field volumes (5x) and potential destructive interference in the scalp caused by reflections oa the skull for 500 kHz (λ = 3 mm) but not for 250 kHz (λ = 6 mm) where wavelength more closely matches scalp thickness. However, where these factors can be controlled, for example during ultrasound of the fingertip, lower frequencies are also more effective in eliciting sensations^34–37,42,51,51,52^. While we cannot assert whether the primary or secondary characteristics of fundamental frequency drive our results, there undoubtedly remains a practical advantage of higher frequencies for confound mitigation. Importantly, there remains a dose-response effect at 500 kHz, highlighting that increasing frequency is not a one-stop solution for somatosensory confounds.

### 4.4. Particle displacement as a primary biophysical driving force underlying peripheral somatosensation

The systematic parameter optimisation approach taken here presents a valuable opportunity to infer the primary biophysical effects that drive neuromodulatory efficacy by leveraging known parameter-biophysics relationships. This approach is one of the few viable methods for making such inferences in healthy human populations. However, limited conclusions can be drawn based on this study alone, and it remains an open question whether peripheral biophysical parameter-effect relationships will translate to the central nervous system.

Putative biophysical mechanisms include acoustic cavitation, particle displacement, acoustic radiation force (ARF), and their respective strain. Cavitation is an unlikely mechanism, as somatosensory co-stimulation occurred well below the cavitation threshold, and empirically observed cavitation is not related to evoked sensations during ultrasound directly focused at the PNS^34^. ARF is dependent on absorption and scales with *f*_0_ and intensity^70^. However, we observed effects that were inversely related to *f*_0_ and scaled linearly with pressure. The observation of stronger effects at lower fundamental frequencies that scale with pressure implicate particle displacement over ARF as the primary driving force behind peripheral somatosensory co-stimulation, in line with findings from peripherally targeted ultrasound^34–36^.

This preliminary evidence for particle displacement as a primary biophysical mechanism does not preclude a complementary role of ARF (strain). In fact, ARF may particularly contribute to tactile sensations, which were most pronounced at a higher ‘dose sharpness’ and in absence of ramping. Here, the sharper (temporal gradient of) ARF displacement could resemble a light ‘tap’ that peripheral mechanoreceptors are highly sensitive to. Nonetheless, the increase in sensations with longer pulse durations across all modalities indicates that a temporally stable component – either sustained ARF displacement or, more likely, the sign-alternating ultrasonic stimulus itself – also contributes to these effects.

It is likely that multiple biophysical effects of ultrasound work in tandem to drive (peripheral) neuromodulation. Future parametric studies can help us converge on a unified theory of key biophysical mechanisms. This pursuit will be critical to identify the principal biophysical effects in PNS and CNS neuromodulation, thus allowing for optimisation of TUS efficacy in the CNS while minimising effects on the PNS. For instance, if ARF were ultimately identified as a central CNS mechanism, as has been suggested^34,45,85,86^, then adaptation towards higher sub-MHz frequencies and ARF interference setups^87^ would become strong avenues to maximise effective dose.

### 4.5. Limitations

This study deliberately applied TUS in a manner expected to cause stronger somatosensory co-stimulation to avoid floor effects and have sufficient sensitivity to detect the effects of changes in stimulation parameters. Specifically, we used lower frequencies (250 kHz), included near-field intensity peaks, and stimulated through the temporal window where somatosensory co-stimulation is more pronounced. Additionally, participants focused on co-stimulation, rather than on a cognitive task that might have reduced confound salience, though this does not negate risks of cueing or ineffective blinding. By operating under conditions that amplify confounds, we reliably mapped parameter-confound relationships, thereby providing actionable strategies to minimise co-stimulation that will also hold at lower confound levels.

Furthermore, we did not directly assess the efficacy of (in)active control conditions or alternative interventions such as topical anaesthetic in blinding participants to stimulation. The latter is unlikely to fully ameliorate (painful) somatosensory co-stimulation considering its primary effects on C-fibres^37^ and its limited efficacy to this end for transcranial electric stimulation^88^. Nevertheless, further research is needed to empirically support optimal controls for somatosensory confounds when present.

### 4.6. Somatosensory confound mitigation strategies

We propose the following workflow to minimise and control for somatosensory confounds in human TUS research. First, the likelihood of peripheral confounds should be assessed during study piloting. If somatosensory co-stimulation is likely, researchers can determine whether transducer-specific characteristics like near-field intensity peaks and energy dispersion in the scalp can be adapted to circumvent confounds. These interventions will have little-to-no impact on CNS neuromodulatory efficacy. Next, pulsing parameters can be optimised by introducing ramping and decreasing ‘dose sharpness’. If somatosensory confounds persist, researchers can consider adjusting fundamental frequency, PRF, or dose itself (Fig. 6). However, undesired impact on CNS neuromodulation should be carefully considered for these manipulations. For example, fundamental frequency selection should holistically balance the required target spatial resolution, as well as the relevant safety metric boundaries, desired primary biophysical effects, and practical constraints^59,70,71^.

**Fig. 6.**
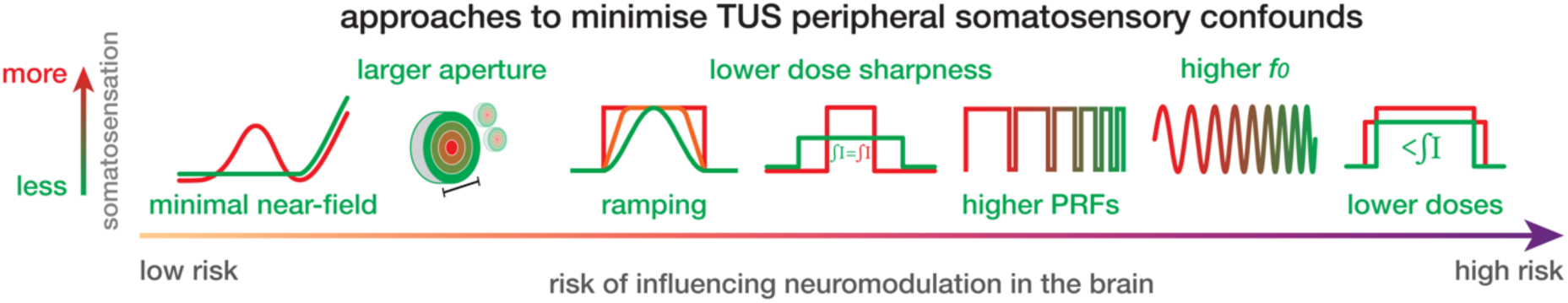
Approaches to minimise TUS peripheral confounds in order of risk of influencing neuromodulation in the brain. Conditions under which somatosensory co-stimulation is less pronounced are illustrated in green.

Robust control conditions will be required in cases where the somatosensory confound cannot be fully alleviated. Common sound-only sham conditions will not sufficiently mimic somatosensory co-stimulation. Therefore, active or inactive control stimulation sites are preferred, where this limitation is addressed by precisely replicating auditory and somatosensory confounds without delivering effective dose. Defocusing the transducer may also be an effective control technique, though this is not possible for all transducers. Furthermore, care should be taken that there is a similar intensity profile in the scalp during verum and control conditions. Using these controls, we can make substantiated inferences on direct neuromodulatory effects, even when peripheral confounds are present.

## Conclusion

Managing somatosensory confounds is critical to minimise participant burden and ensure valid and replicable findings as TUS research progresses toward higher doses, more frequent transducer placement at the sensitive temples of the head, and smaller transducers. This study characterises the range of somatosensory co-stimulation experienced during TUS and identifies effective mitigation strategies. These include reducing near-field intensity peaks in the scalp, dispersing energy across the scalp, ramping the pulse envelope, and lowering ‘dose sharpness’. Higher pulse repetition frequencies, higher fundamental frequencies, and lower doses further minimise these effects. Where confounds cannot be fully resolved, robust control conditions, such as (in)active controls that replicate auditory and somatosensory confounds, are essential to isolate direct neuromodulatory effects. By adopting these strategies, researchers can enhance the reliability of TUS research and accelerate effective ultrasonic neuromodulation in scientific, commercial, and clinical domains.

## Data availability

Data and code to reproduce the results reported in this study will be made available following peer review.

## CRediT authorship contribution statement

**Benjamin R. Kop**: conceptualisation, methodology, software, validation, formal analysis, investigation, data curation, writing – original draft, writing – review & editing, visualisation, supervision, project administration.

**Linda de Jong:** methodology, investigation, writing – review & editing

**Kim Butts Pauly:** writing – review & editing

**Hanneke E.M. den Ouden:** writing – review & editing, supervision, funding acquisition

**Lennart Verhagen:** conceptualisation, resources, writing – review & editing, supervision, funding acquisition

## Acknowledgements

We thank Soha Farboud, Sjoerd Meijer, and Dr. Marijtje Jongsma for our discussions. We also thank Margely Cornelissen and Stein Fekkes from the Radboud FUS Initiative. LV is supported by the Dutch Research Council (NWO, VIDI fellowship, 18919), the European Innovation Council (EIC, co-applicant Pathfinder project CITRUS, 101071008), and the European Research Council (ERC, co-applicant MediCoDe). HEMdO is supported by the Dutch Research Council (NWO, VIDI fellowship, 175.450).

## Funding

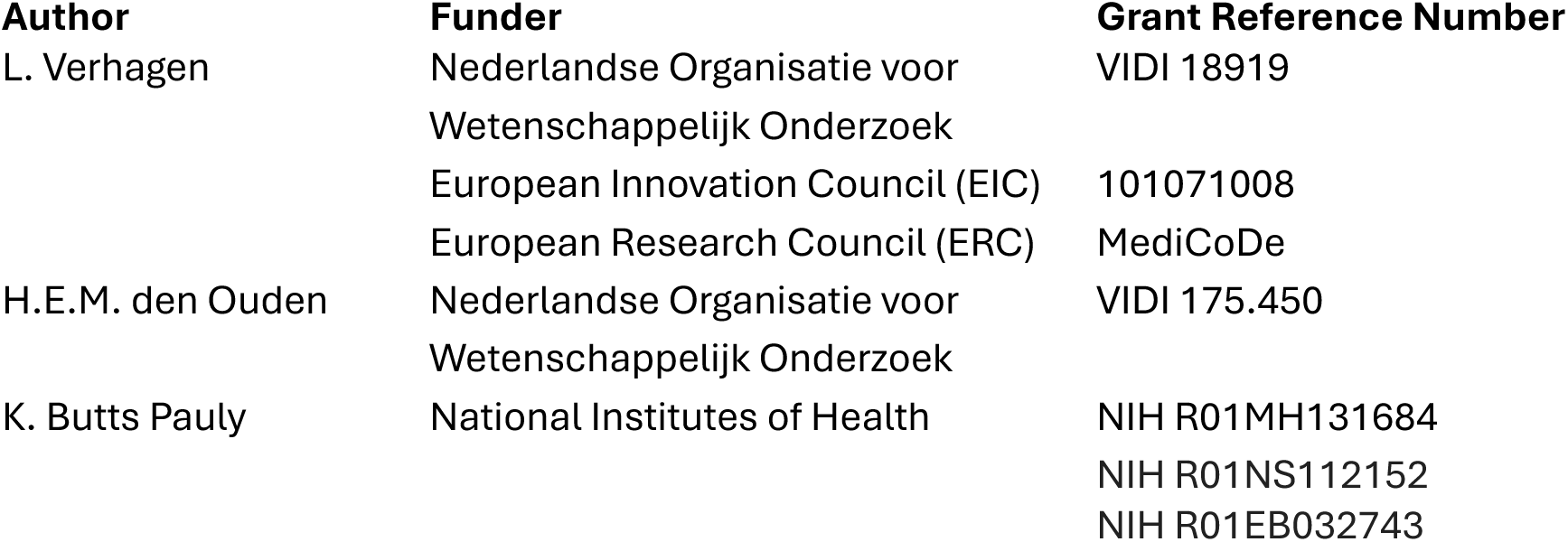

## A Supplementary Material

**Supplementary Table 1.**
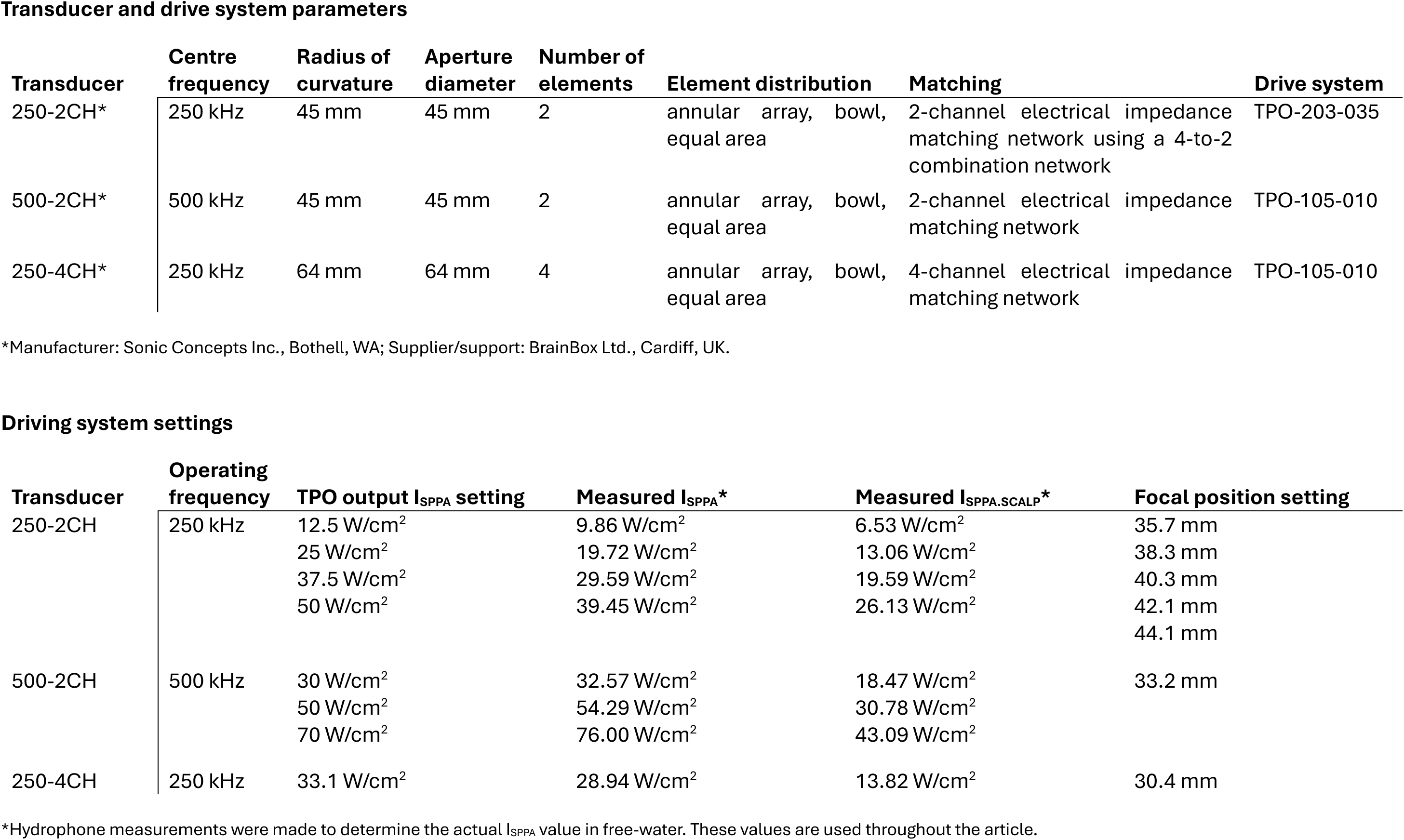
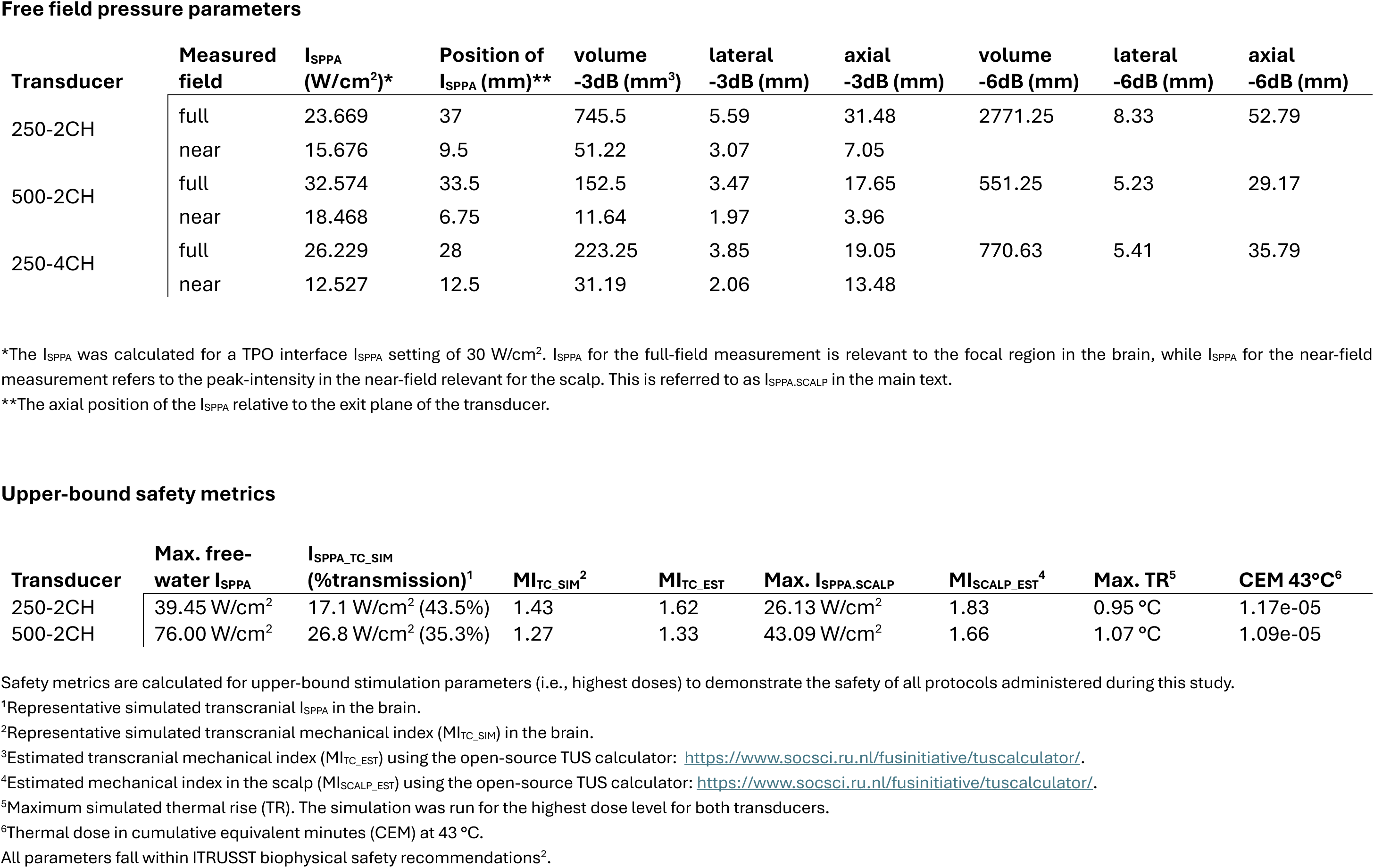
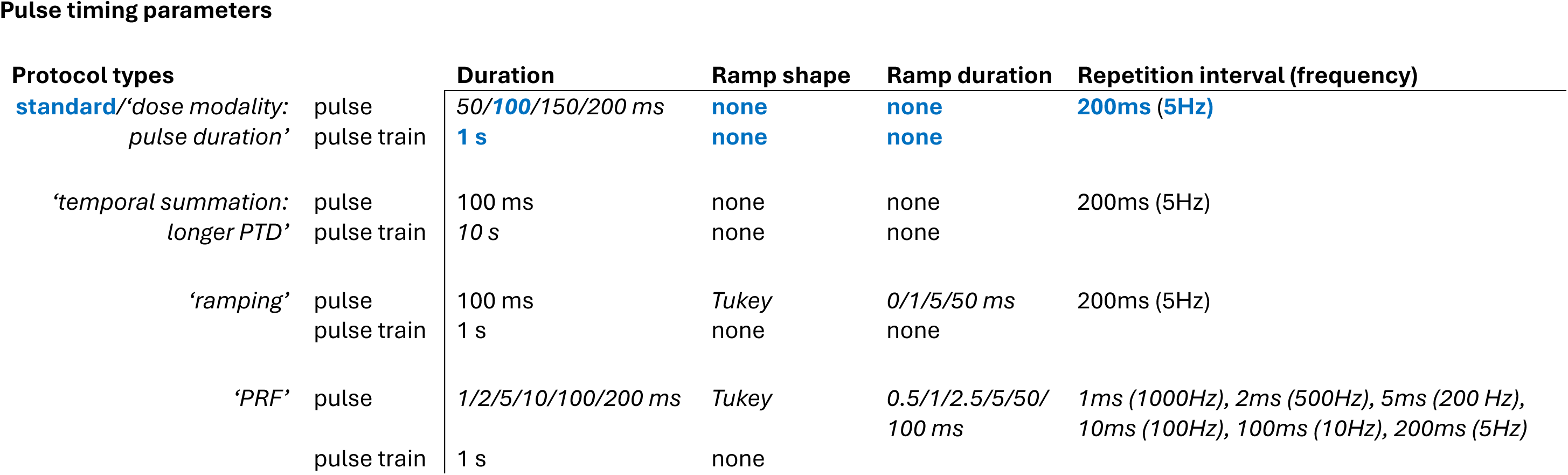
TUS specifications per ITRUSST standardised reporting^1^ guidelines. This table depicts the pulse timing parameters for the standard protocol in blue, as well as the relevant timing for manipulated parameters in italics. This includes the investigation of dose by varying pulse duration, the longer pulse train duration (PTD) applied to mimic offline protocols while participants continuously rated somatosensory co-stimulation, ramping, and pulse repetition frequency (PRF) where full ramping was administered.

**Transducer and drive system parameters**
| Transducer | Centre frequency | Radius of curvature | Aperture diameter | Number of elements | Element distribution | Matching | Drive system |
| --- | --- | --- | --- | --- | --- | --- | --- |
| 250-2CH* | 250 kHz | 45 mm | 45 mm | 2 | annular array, bowl, equal area | 2-channel electrical impedance matching network using a 4-to-2 combination network | TPO-203-035 |
| 500-2CH* | 500 kHz | 45 mm | 45 mm | 2 | annular array, bowl, equal area | 2-channel electrical impedance matching network | TPO-105-010 |
| 250-4CH* | 250 kHz | 64 mm | 64 mm | 4 | annular array, bowl, equal area | 4-channel electrical impedance matching network | TPO-105-010 |

\*Manufacturer: Sonic Concepts Inc., Bothell, WA; Supplier/support: BrainBox Ltd., Cardiff, UK.

**Driving system settings**
| Transducer | Operating frequency | TPO output I<sub>SPPA</sub> setting | Measured I<sub>SPPA</sub>* | Measured I<sub>SPPA.SCALP</sub>* | Focal position setting |
| --- | --- | --- | --- | --- | --- |
| 250-2CH | 250 kHz | 12.5 W/cm<sup>2</sup> | 9.86 W/cm<sup>2</sup> | 6.53 W/cm<sup>2</sup> | 35.7 mm |
|  |  | 25 W/cm<sup>2</sup> | 19.72 W/cm<sup>2</sup> | 13.06 W/cm<sup>2</sup> | 38.3 mm |
|  |  | 37.5 W/cm<sup>2</sup> | 29.59 W/cm<sup>2</sup> | 19.59 W/cm<sup>2</sup> | 40.3 mm |
|  |  | 50 W/cm<sup>2</sup> | 39.45 W/cm<sup>2</sup> | 26.13 W/cm<sup>2</sup> | 42.1 mm |
|  |  |  |  |  | 44.1 mm |
| 500-2CH | 500 kHz | 30 W/cm<sup>2</sup> | 32.57 W/cm<sup>2</sup> | 18.47 W/cm<sup>2</sup> | 33.2 mm |
|  |  | 50 W/cm<sup>2</sup> | 54.29 W/cm<sup>2</sup> | 30.78 W/cm<sup>2</sup> |  |
|  |  | 70 W/cm<sup>2</sup> | 76.00 W/cm<sup>2</sup> | 43.09 W/cm<sup>2</sup> |  |
| 250-4CH | 250 kHz | 33.1 W/cm<sup>2</sup> | 28.94 W/cm<sup>2</sup> | 13.82 W/cm<sup>2</sup> | 30.4 mm |

\*Hydrophone measurements were made to determine the actual I<sub>SPPA</sub> value in free-water. These values are used throughout the article.

Free field pressure parameters
| Transducer | Measured field | $I_{SPPA}$ (W/cm<sup>2</sup>)* | Position of $I_{SPPA}$ (mm)** | volume -3dB (mm<sup>3</sup>) | lateral -3dB (mm) | axial -3dB (mm) | volume -6dB (mm) | lateral -6dB (mm) | axial -6dB (mm) |
| --- | --- | --- | --- | --- | --- | --- | --- | --- | --- |
| 250-2CH | full | 23.669 | 37 | 745.5 | 5.59 | 31.48 | 2771.25 | 8.33 | 52.79 |
|  | near | 15.676 | 9.5 | 51.22 | 3.07 | 7.05 |  |  |  |
| 500-2CH | full | 32.574 | 33.5 | 152.5 | 3.47 | 17.65 | 551.25 | 5.23 | 29.17 |
|  | near | 18.468 | 6.75 | 11.64 | 1.97 | 3.96 |  |  |  |
| 250-4CH | full | 26.229 | 28 | 223.25 | 3.85 | 19.05 | 770.63 | 5.41 | 35.79 |
|  | near | 12.527 | 12.5 | 31.19 | 2.06 | 13.48 |  |  |  |

\*The $I_{SPPA}$ was calculated for a TPO interface $I_{SPPA}$ setting of 30 W/cm<sup>2</sup>. $I_{SPPA}$ for the full-field measurement is relevant to the focal region in the brain, while $I_{SPPA}$ for the near-field measurement refers to the peak-intensity in the near-field relevant for the scalp. This is referred to as $I_{SPPA,SCALP}$ in the main text.
\*\*The axial position of the $I_{SPPA}$ relative to the exit plane of the transducer.

Upper-bound safety metrics
| Transducer | Max. free-water $I_{SPPA}$ | $I_{SPPA,TC\_SIM}$ (%transmission)<sup>1</sup> | $MI_{TC\_SIM}$<sup>2</sup> | $MI_{TC\_EST}$ | Max. $I_{SPPA,SCALP}$ | $MI_{SCALP\_EST}$<sup>4</sup> | Max. TR<sup>5</sup> | CEM 43°C<sup>6</sup> |
| --- | --- | --- | --- | --- | --- | --- | --- | --- |
| 250-2CH | 39.45 W/cm<sup>2</sup> | 17.1 W/cm<sup>2</sup> (43.5%) | 1.43 | 1.62 | 26.13 W/cm<sup>2</sup> | 1.83 | 0.95 °C | 1.17e-05 |
| 500-2CH | 76.00 W/cm<sup>2</sup> | 26.8 W/cm<sup>2</sup> (35.3%) | 1.27 | 1.33 | 43.09 W/cm<sup>2</sup> | 1.66 | 1.07 °C | 1.09e-05 |

Pulse timing parameters
| Protocol types |  | Duration | Ramp shape | Ramp duration | Repetition interval (frequency) |
| --- | --- | --- | --- | --- | --- |
| **standard**/'dose modality:<br>pulse duration' | pulse | 50/**100**/150/200 ms | **none** | **none** | **200ms (5Hz)** |
|  | pulse train | **1 s** | **none** | **none** |  |
| 'temporal summation:<br>longer PTD' | pulse | 100 ms | none | none | 200ms (5Hz) |
|  | pulse train | 10 s | none | none |  |
| 'ramping' | pulse | 100 ms | *Tukey* | *0/1/5/50 ms* | 200ms (5Hz) |
|  | pulse train | 1 s | none | none |  |
| 'PRF' | pulse | *1/2/5/10/100/200 ms* | *Tukey* | *0.5/1/2.5/5/50/*<br>*100 ms* | *1ms (1000Hz), 2ms (500Hz), 5ms (200 Hz),*<br>*10ms (100Hz), 100ms (10Hz), 200ms (5Hz)* |
|  | pulse train | 1 s | none |  |  |

**Supplementary Table 2.** MRI acquisition parameters. This table shows the MRI acquisition parameters. Scans were acquired using a 3T Siemens Skyra MRI scanner (Siemens Medical Solutions, Erlangen, Germany) with a 32-channel head coil. Anatomical T1w scans were used for online neuronavigation. UTE scans were acquired to generate pseudo-CT images for post-hoc simulation

| Scan | TR | TE | FoV read | FoV phase | voxel | N slices |
| --- | --- | --- | --- | --- | --- | --- |
| T1w | 2700 ms | 3.69 ms | 230 mm | 128.1% | 0.9 mm iso | 224 |
| UTE | 3.6 ms | 0.07 ms | 240 mm | 100.0% | 0.8 mm iso | 320 |

**Supplementary Table 3.**
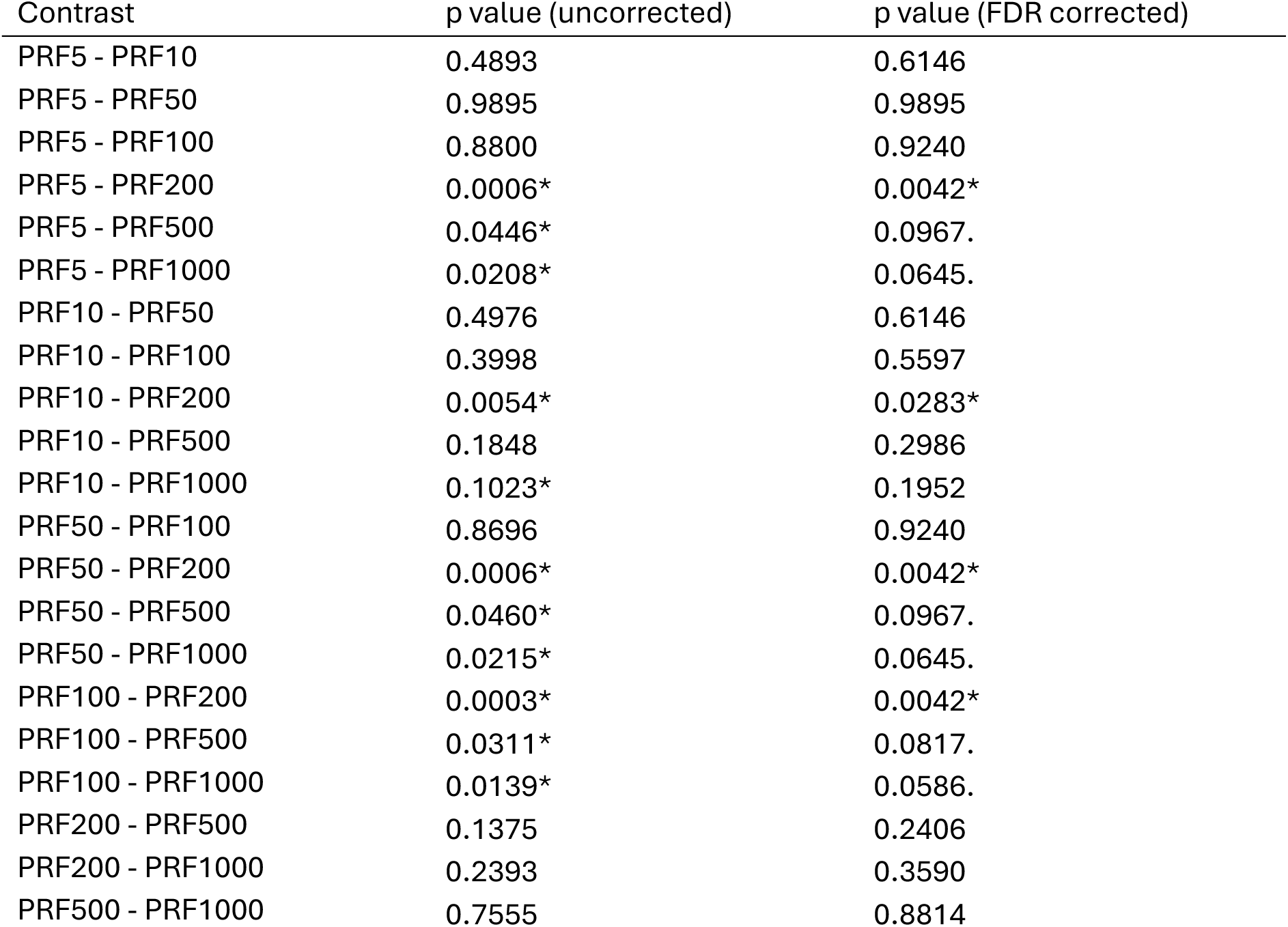
Post-hoc pairwise comparisons for the eDect of PRF on thresholds. Post-hoc paired comparisons between each applied level of PRF, both without correction for multiple comparison (middle column), and with false discovery rate (FDR) correction for multiple comparisons. ‘*’ = significant, ‘.’ = trend.

| Contrast | p value (uncorrected) | p value (FDR corrected) |
| --- | --- | --- |
| PRF5 - PRF10 | 0.4893 | 0.6146 |
| PRF5 - PRF50 | 0.9895 | 0.9895 |
| PRF5 - PRF100 | 0.8800 | 0.9240 |
| PRF5 - PRF200 | 0.0006* | 0.0042* |
| PRF5 - PRF500 | 0.0446* | 0.0967. |
| PRF5 - PRF1000 | 0.0208* | 0.0645. |
| PRF10 - PRF50 | 0.4976 | 0.6146 |
| PRF10 - PRF100 | 0.3998 | 0.5597 |
| PRF10 - PRF200 | 0.0054* | 0.0283* |
| PRF10 - PRF500 | 0.1848 | 0.2986 |
| PRF10 - PRF1000 | 0.1023* | 0.1952 |
| PRF50 - PRF100 | 0.8696 | 0.9240 |
| PRF50 - PRF200 | 0.0006* | 0.0042* |
| PRF50 - PRF500 | 0.0460* | 0.0967. |
| PRF50 - PRF1000 | 0.0215* | 0.0645. |
| PRF100 - PRF200 | 0.0003* | 0.0042* |
| PRF100 - PRF500 | 0.0311* | 0.0817. |
| PRF100 - PRF1000 | 0.0139* | 0.0586. |
| PRF200 - PRF500 | 0.1375 | 0.2406 |
| PRF200 - PRF1000 | 0.2393 | 0.3590 |
| PRF500 - PRF1000 | 0.7555 | 0.8814 |

**Supplementary Fig. 1.**
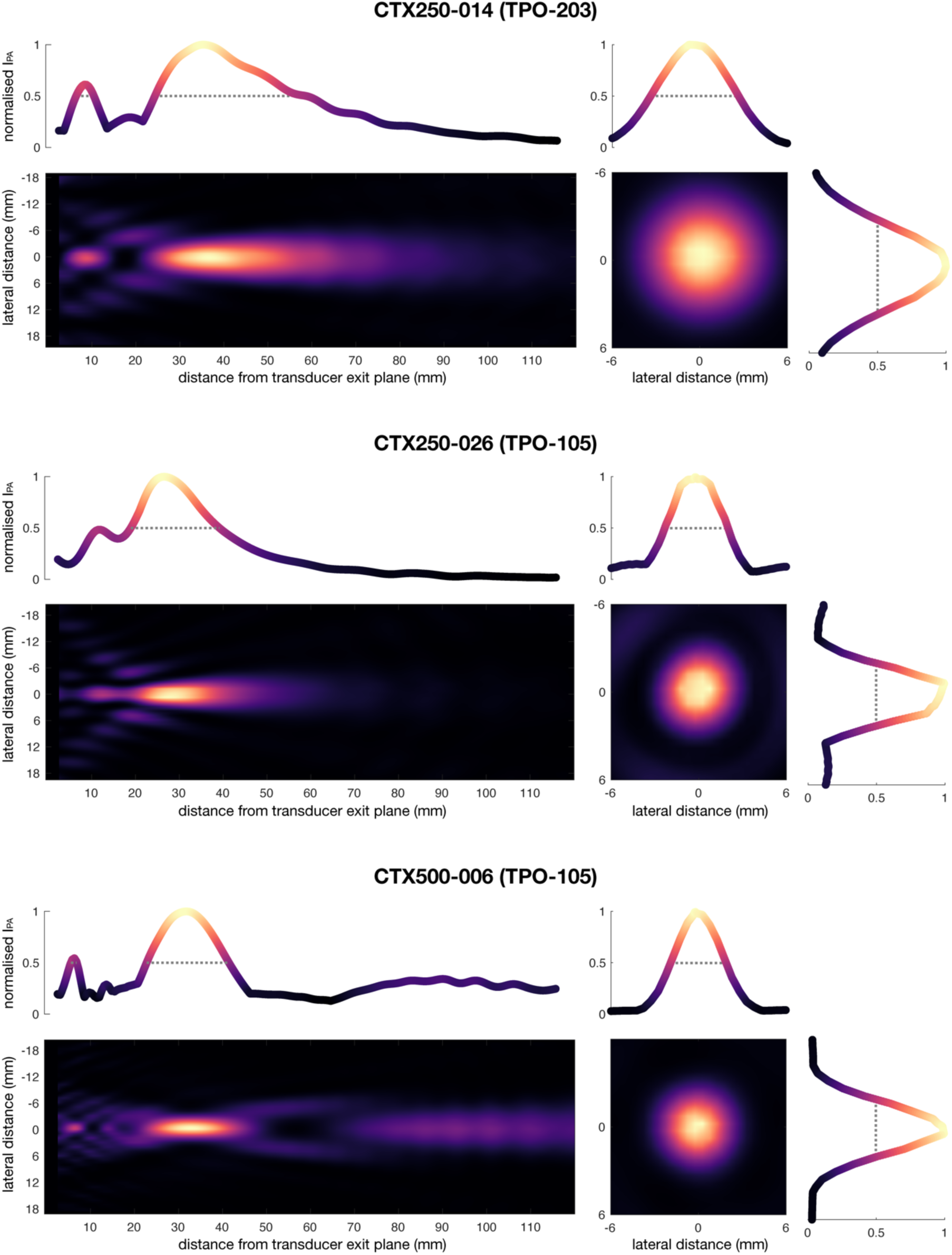
Full-field hydrophone measurements. Hydrophone measurements of the min-max normalised pulse-average intensity (IPA) for the full acoustic field along the axial plane (left) and the lateral cross-section at the focus (right) of each transducer. The original resolution (0.5 mm) has been upscaled for visualisation by a factor of 10 using linear interpolation. Intensity distribution lines depict the maximum IPA per slice, where the full-width-half-maximum of the ISPPA (FWHM; -3dB) is indicated by the dotted grey line.

**Supplementary Fig. 2.**
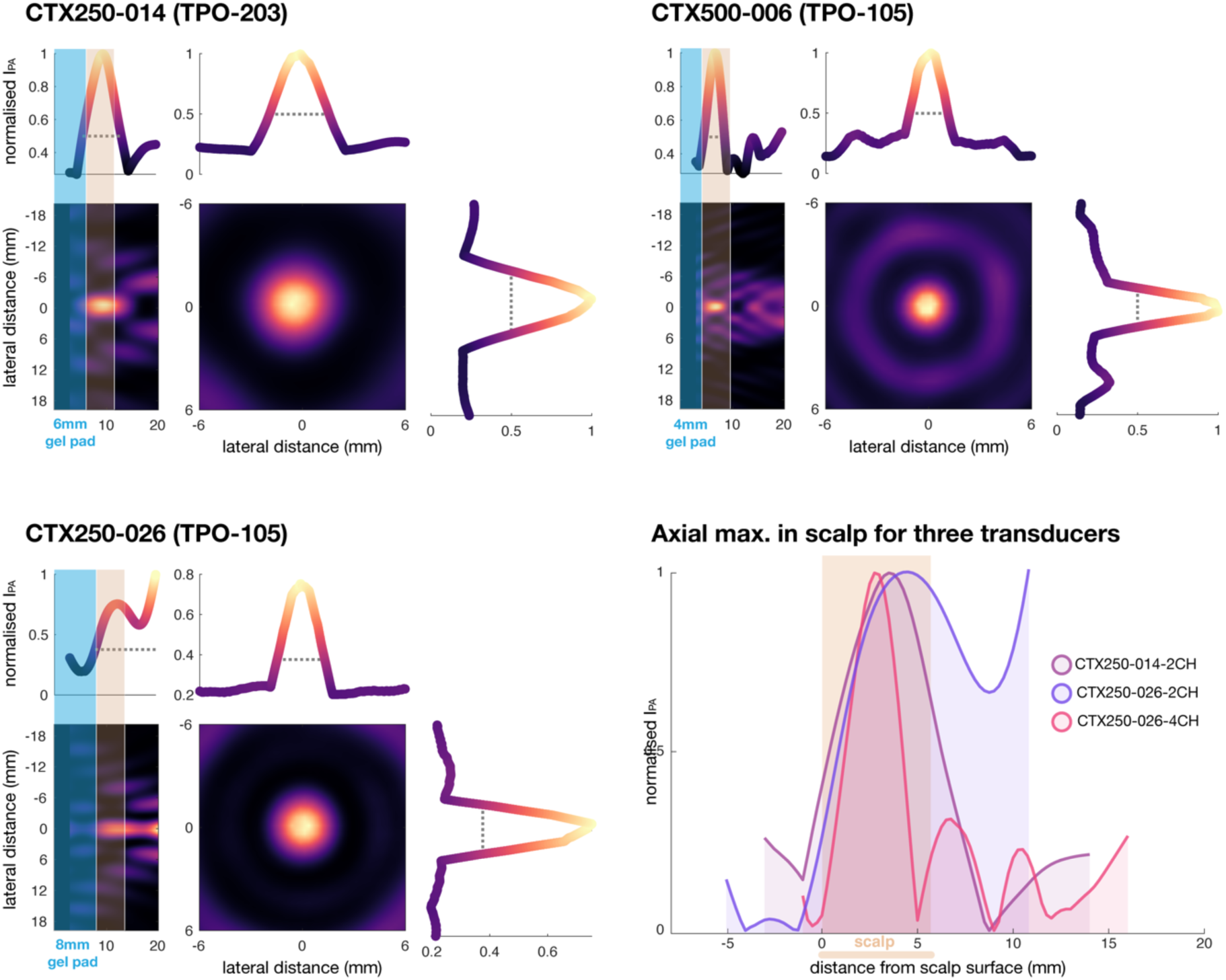
Near-field hydrophone measurements. Hydrophone measurements of the min-max normalised pulse-average intensity (IPA) for the near-field at an increased resolution of 0.25 mm. These data were used to determine the gel pad thicknesses (blue) and stimulation intensities. Note that the bottom-right panel depicts the normalised distributions of the axial maximum intensity for each transducer. In practice, the relative stimulation intensities of the transducers were adjusted to achieve comparable levels of integrated maximum intensity and integrated total intensity in the scalp (beige) between transducers (see main text Fig. 4).

Near-field hydrophone measurements were required to accurately assess the effects of fundamental frequency and transducer aperture diameter on peripheral somatosensory co-stimulation. These investigations involved different transducers with varying intensity profiles, which had to be equalised to make valid comparisons.

To this end, we utilised the integrated maximum and/or total intensity in the scalp (5.5 mm width^3–5^) as a metric to optimise comparability between transducers. First, we identified focal depth settings at which the acoustic profiles were most similar (250-2CH: 40.3 mm; 250-4CH: 30.4 mm; 500-2CH: 33.1 mm). Next, we determined the gel pad thicknesses to optimise coherence of acoustic profiles and the integrated maximum and/or total intensity in the scalp (250-2CH: 6 mm; 250-4CH: 8 mm; 500-2CH: 4 mm). Finally, we set stimulation intensities in the scalp. For equal integrated maximum intensities, we compared fundamental frequencies using transducer 250-2CH at 26.13 W/cm^2^ and transducer 500-2CH at 30.78 W/cm^2^ I_SPPA.SCALP_. For equal integrated total intensity, the intensity for 500-2CH was increased to 43.09 W/cm^2^. The intensities used to compare transducer aperture diameter between 250-2CH and 250-4CH were 13.06 W/cm^2^ and 13.82 W/cm^2^ I_SPPA.SCALP_, respectively.

Hydrophone measurements were performed using an independent metrology setup enabling accurate positioning of a calibrated hydrophone (d_x,y,z_ = 5 µm; HGL 0200, Onda Corp., Sunnyvale, USA). The transducer was submerged in degassed, filtered, and deionised water at ambient temperature in a plexiglass water tank (150x200x400 mm). A custom probe holder ensured orthogonal alignment of the transducer and hydrophone.

The transducer was set to deliver 250 µs square-wave pulses at a power of 2.5 or 5.0 W per channel. These pulses were registered using a PicoScope 5244D (Pico Technology, UK) at a sampling frequency of 25 MHz using a custom closed-loop control program triggered by the transducer power output system.

For full-field measurements, line scans were performed with 0.5 mm steps along the beam axis, centred on the focus, at distances from 3 to 120 mm relative to the exit plane of the transducer. A ∼38 mm range was measured across the lateral cross-sections of the ultrasound beam. Full-field measurements were acquired to inform transcranial ultrasound for the experiment.

To capture a higher resolution intensity field for depths relevant to peripheral stimulation of the scalp, we recorded near-field intensities using 0.25 mm steps for an axial range of 3 to 20 mm from the transducer exit plane, and a lateral range of ∼40 mm. Near-field measurements were acquired to inform the gel pad thicknesses and absolute free-water intensities required to equalise the integrated maximum intensity and/or integrated total intensity in the scalp.

Post-processing involved an FFT-based method to acquire a single average amplitude reading for each recorded pulse over the window between the pulse ring-up time and pulse cessation. Subsequently, the complete pressure field was spatially filtered along the axial direction using a FIR Butterworth low-pass filter to remove oscillating interference caused by reflections off the hydrophone.

The measured intensities were then re-scaled using the factor Power_experiment_/Power_hydrophone.measurement_. Next, the location and value of the spatial-peak pulse-average intensity at the focus (I_SPPA_) and the peak near-field intensity (I_SPPA.SCALP_) were extracted and the focal dimensions of the -3dB focal region were calculated using in the *regionprops3* MATLAB function.

Both full- and near-field measurements were performed for three transducers (i.e., 250-2CH – TPO-203, 500-2CH – TPO-105, and 250-4CH – TPO-105; see Supplementary Table 2 for detailed specifications), at focal depth settings of 40.3, 33.2, and 30.4 mm, respectively.

**Supplementary Fig. 3.**
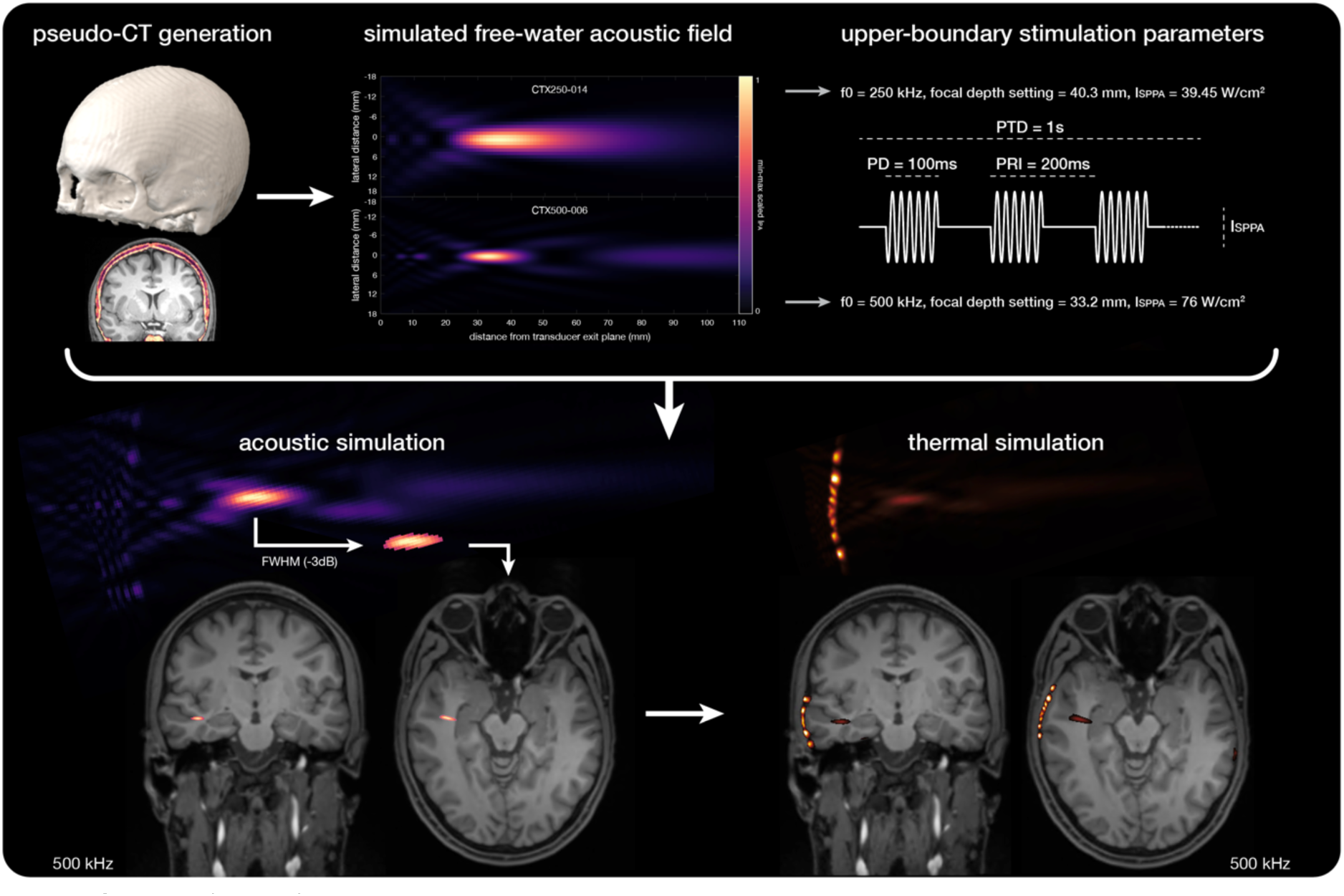
Simulations. Pseudo-CT scans (top left) were generated from UTE scans and used to assign acoustic medium properties. Next, we confirmed that the free-field simulations for both 250-2CH and 500-2CH transducers corresponded well with hydrophone measurements. We then ran acoustic and thermal simulations for upper-bound stimulation parameters to obtain safety-relevant metrics (top right). The transcranial full-width half-maximum (FWHM) intensity for 500 kHz stimulation is depicted on the bottom left. The subsequent thermal simulation is depicted on the right.

We ran a representative simulation of acoustic wave propagation for 250 kHz and 500 kHz TUS using k-Plan, a user interface for the pseudo-spectral time-domain solver k-Wave^6^. First, we generated a pseudo-CT (pCT) scan from our ultra-short echo time (UTE) MRI scan using the open-source ‘petra-to-pct’ toolbox (https://github.com/ucl-bug/petra-to-ct)^7^. Histogram normalisation was set to two peaks at a minimum distance of 1000 units and skull mask smoothing was set to 5 mm.

In k-Plan, we first simulated our custom transducer models in free-water and confirmed that the full-field intensity profile was comparable to our hydrophone measurements. Next, we ran acoustic and thermal simulations to assess acoustic targeting and to estimate upper-boundary safety metrics for these transducers. The maximum dose was simulated with: PD = 100 ms, PRI = 200 ms, PTD = 1 s, ISPPA.FREE.WATER = 39.45 W/cm^2^ (250 kHz) or 76 W/cm^2^ (500 kHz).

The simulated pressure field was exported using the ‘k-plan-matlab-tools’ toolbox (https://github.com/ucl-bug/k-plan-matlab-tools). In MATLAB, the intensity field was calculated using 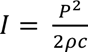, where ρ was 1000 kg/m^3^ and *c* was 1500 m/s. A full-width half-maximum threshold (-3dB) was then applied to the intensity field, and the resulting field was overlaid onto the MRI scan.

**Supplementary Fig. 4.**
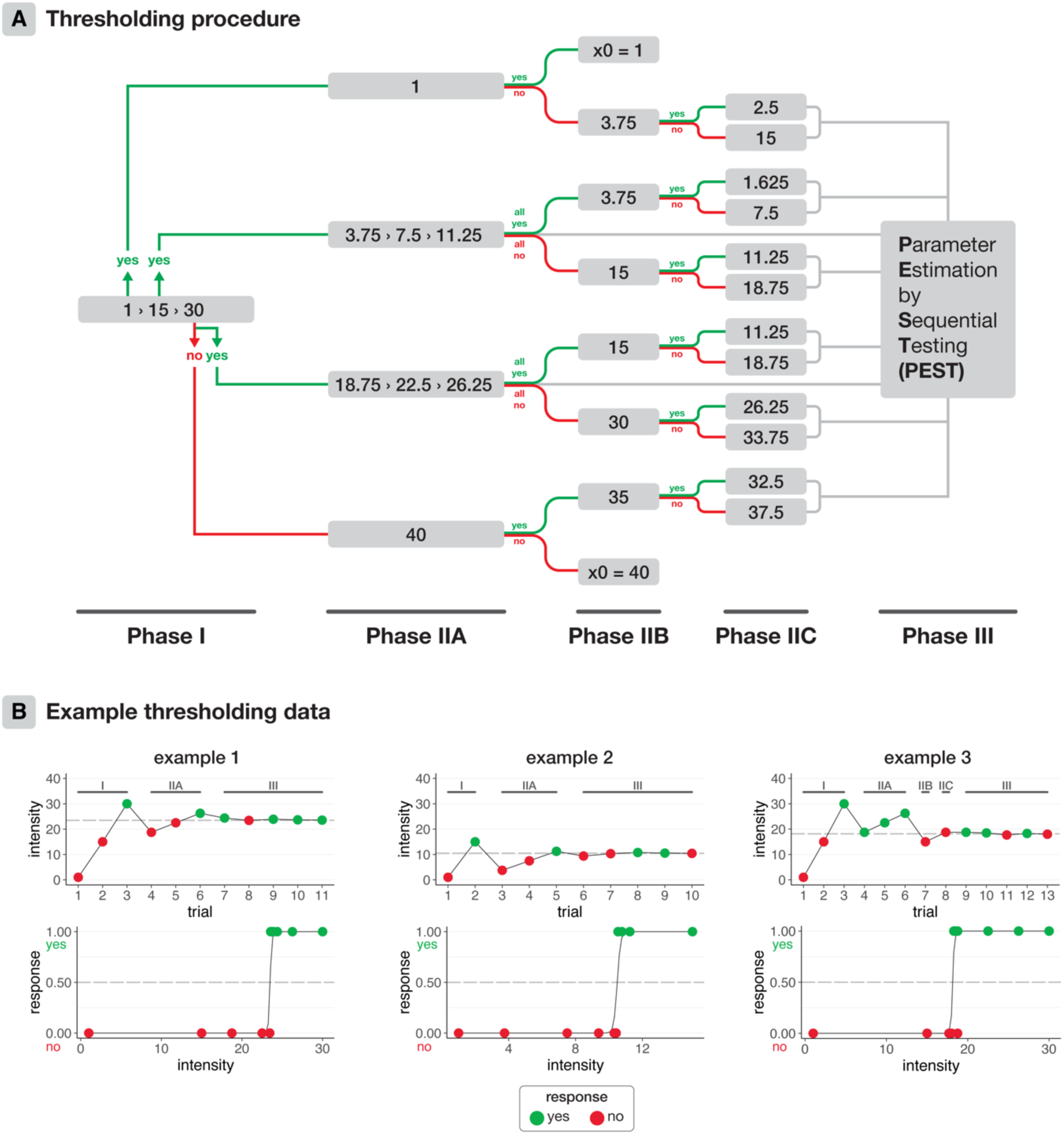
Thresholding procedure. **(A)** fiowchart of the custom thresholding procedure. **(B)** Example thresholding data. For the top panels, the TPO interface ISPPA is displayed across trials to demonstrate the operation of the thresholding procedure. Green and red dots indicate ‘yes’ and ‘no’ responses respectively to whether the stimulus was felt. The dotted grey line depicts the estimated sensory threshold. The bottom panel depicts the fitted psychometric curve over the binary yes/no responses after completion of the thresholding procedure. In examples 1 and 2, the participant transitioned between ‘no’ and ‘yes’ within the three intermediate stimulus intensities tested in Phase IIA, therefore continuing directly to Parameter Estimation by Sequential Testing (PEST; Phase III). In example 3, the participant responded yes to all three intermediate intensities, so the lower boundary was re-tested (Phase IIB). Since the response was ‘no’, a slightly higher boundary was tested (Phase IIC). Then, the thresholding procedure continued to Phase III.

We measured sensory thresholds to precisely capture the effects of pulse repetition frequency and ramping on somatosensory co-stimulation. Typically, many trials are required to estimate sensory thresholds^8^. In the present experiment, that would have required a prohibitively large amount of ultrasonic stimulation. Therefore, we designed a custom thresholding procedure that consisted of three phases (Supplementary Fig. 4A).

Phase III consisted of five trials where we iteratively fit a logistic, psychometric function to the binary response data of all preceding trials using a Parameter Estimation by Sequential Testing (PEST) method^9,10^. We defined the logistic function as:

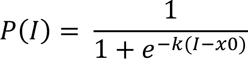

where *I* is the TPO interface I_SPPA_ value, *x0* is the stimulus intensity at which the detection probability is an estimated 50%, and *k* is the slope of the psychometric curve. The curve was fit using the ‘curve_fit’ function from the SciPy package in Python. Initial parameter values for *x0* and *k* were set to the *t-1* stimulus intensity and 1, respectively, to improve convergence. We constrained the optimisation with bounds of -5-40 for *x0* and 0.01-100 for *k*.

In some cases, participants already reported feeling stimulation at a TPO intensity setting of 1 W/cm^2^, or didn’t report feeling anything from 1-30 W/cm^2^. In the prior case, Phase IIA re-tested this minimum stimulus intensity. If the participant continued to respond ‘yes’, 1 W/cm^2^ was set as their threshold (floor effect). When participants did not report feeling anything from 1-30 W/cm^2^, in Phase IIA stimulation intensity was increased to 40 W/cm^2^. If participants still did not feel anything, 40 W/cm^2^ was set as their threshold (ceiling effect).

**Supplementary Fig. 5.**
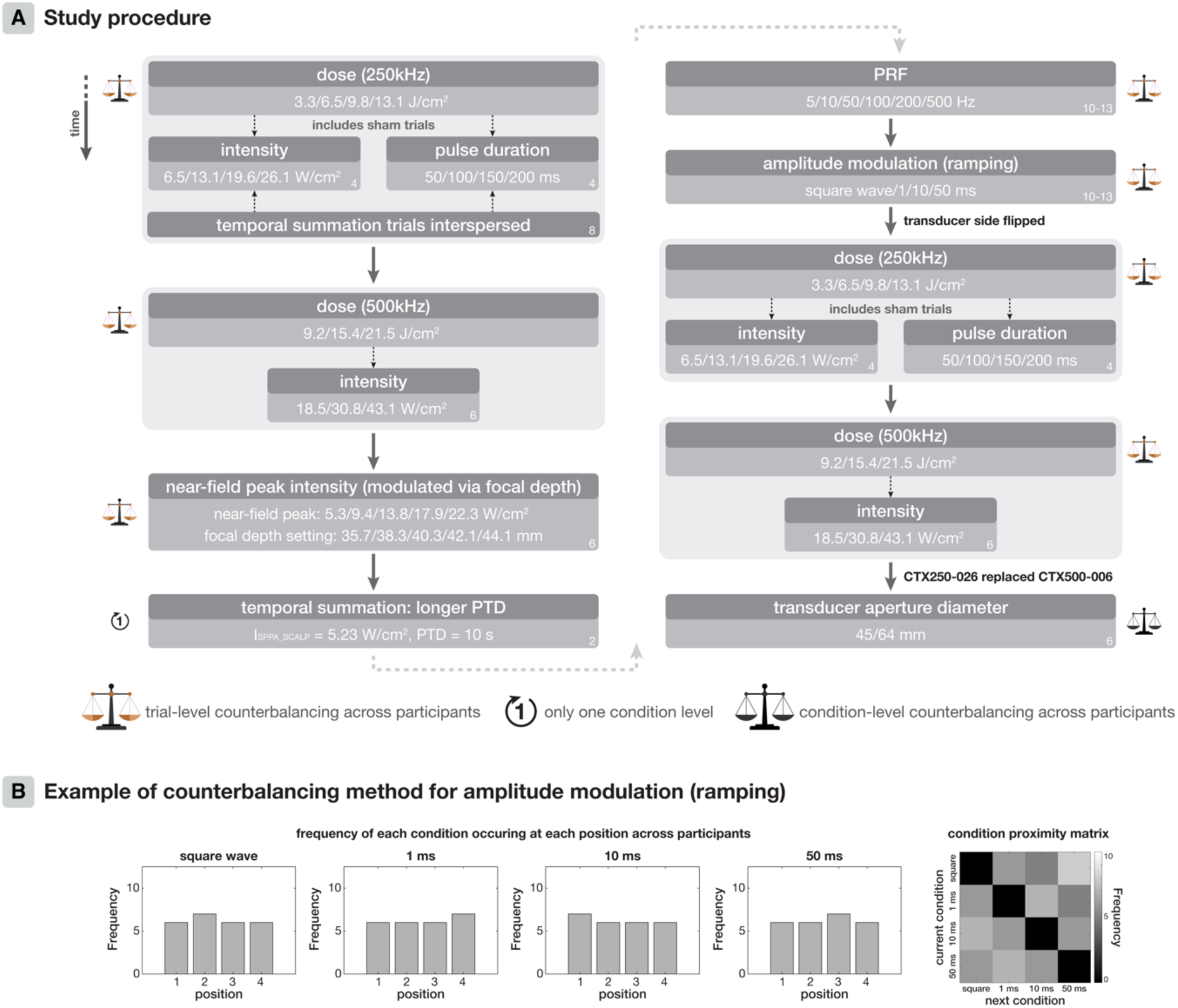
Study procedure and counterbalancing **(A)** Study procedure depicting the order of investigations taking place during the main experiment, with the type of counterbalancing indicated. **(B)** Counterbalancing example. We minimised variance in the frequency of conditions occurring across trial positions and the direct succession of conditions in one pair occurring more often than another pair. The frequency represents the number of participants in which that condition (e.g., square wave) occurs at each trial position.

At the beginning of the experiment, participants completed practice trials, including the lowest and highest doses for both transducers (250-2CH & 500-2CH), to familiarise themselves with VAS ratings and get an idea of what to expect during the experiment.

Next, different TUS parameters were manipulated in the order specified in Supplementary Fig. 5A. This uniform order was used so that any cumulative effects of stimulation across the experiment were equal for each investigated parameter, such that the total energy applied prior to each separate ‘research question’ was the same. To account for any possible interaction between temporal summation of peripheral somatosensation over time and differences in the initial side that 250 kHz and 500 kHz stimulation were applied, the starting side of the two transducers was also counterbalanced across participants.

Participants took a short break after the ‘temporal summation: longer PTD’ and ‘amplitude modulation (ramping)’ segments. Transducers were re-positioned and re-coupled at these times, as well as prior to the ‘transducer aperture diameter’ segment. Throughout the experiment, coupling quality was visually monitored at ∼5-minute intervals.

For all manipulated stimulation parameters except for ‘transducer aperture diameter’, we implemented trial-level counterbalancing across participants. First, we generated all possible orders of unique condition levels using MATLAB 2019b for each of the following parameters: intensity (250 kHz), pulse duration (250 kHz), intensity (500 kHz), near-field peak intensity modulated via focal depth, PRF and ramping. We then identified the subset of N = 25 orders that would optimise trial-level counterbalancing per condition.

We evaluated counterbalancing quality via two metrics. First, we determined the frequency of each condition being administered at each trial position across participants, aiming to minimise variability in these frequencies to ensure that conditions were distributed as evenly as possible. Second, we determined how often two specific condition levels occurred consecutively, aiming to prevent specific pairs of conditions from occurring more often than other pairs.

Specifically, we compiled 1e9 random sets of N=25 condition orders for each manipulated parameter. From these, we selected the set with the lowest variance in condition frequency across participants and minimal variance in consecutive condition transitions (Supplementary Fig. 5B).

For VAS measurements, the same condition was repeated multiple times (*n*) per participant (see bottom-right of each box in Supplementary Fig. 5A). Here, the same order of conditions was presented *n* times. While this repetition could potentially amplify condition order effects within individuals, this approach mitigates the risk that temporal summation of somatosensory co-stimulation across trials could differentially impact different conditions within a single participant. Moreover, applying conditions in sets allowed for comparisons between successive sets to assess temporally summative effects across multiple protocols simultaneously (see main text Fig. 4D). By counterbalancing across participants, we have effectively controlled for condition order effects at the between-subject level.

The investigation of dose for 250 kHz stimulation included counterbalanced orders generated separately for ‘intensity’ and ‘pulse duration’ modalities, each including the sham condition as a level. This design resulted in 8 counterbalanced sham trials delivered to each side of the head. The orders for ‘intensity’ and ‘pulse duration’ dose modalities were then interleaved, with the starting modality counterbalanced between participants. Additionally, in the first ‘dose’ block we included interspersed trials with our standard protocol at ∼1 minute intervals to monitor potential temporally summative effects on somatosensory co-stimulation across trials (see Supplementary Fig. 9).

To investigate the effects of transducer aperture diameter, we administered six consecutive trials each of the standard stimulation protocol using the two-element 250-2CH and four-element 250-4CH. Here, conditions were measured consecutively to capture any temporal summation of somatosensory co-stimulation for identical consecutive trials (see Supplementary Fig. 9). Which transducer was tested first was counterbalanced across participants, as was the side of the transducers.

Sensory thresholds were measured for ‘PRF’ and ‘ramping’ sub-experiments. Here, 10-13 trials of the same protocol, administered at different intensities, were repeated to find the intensity at which the participant could perceive the stimulus 50% of the time (see Supplementary Fig. 4 for full details on the thresholding procedure).

Finally, to assess temporal summation of somatosensory co-stimulation during a longer PTD, mimicking the types of protocols applied in ‘offiine’ TUS studies, we administered the standard protocol at an ISPPA.SCALP of 5.23 W/cm^2^ for a 10 second PTD and participants continuously reported their sensations on a VAS. Participants practiced the continuous VAS scale once without TUS and then completed this procedure twice with 10 s PTD TUS.

**Supplementary Fig. 6.**
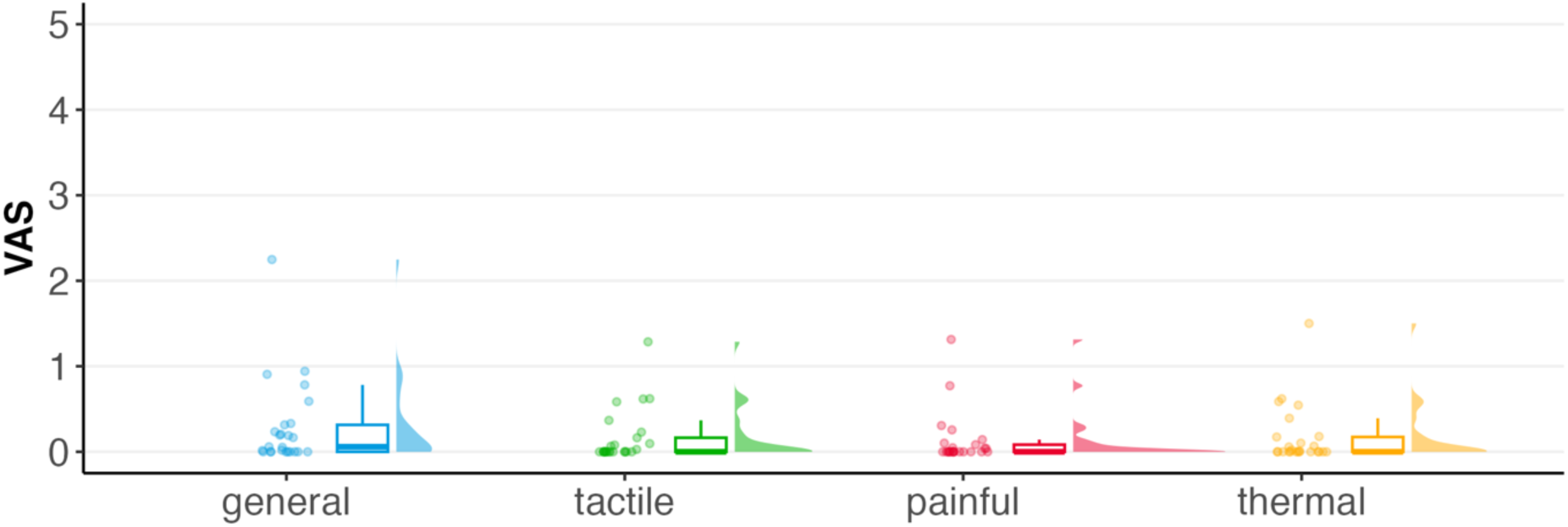
VAS responses to sham trials. The sham condition, consisting of an auditory stimulus administered over speakers, elicited minor somatosensory e@ects in some participants. Points represent participant-level medians used for sham-correction. Boxplots and half-violins reflect the distribution of the data.

**Supplementary Fig. 7.**
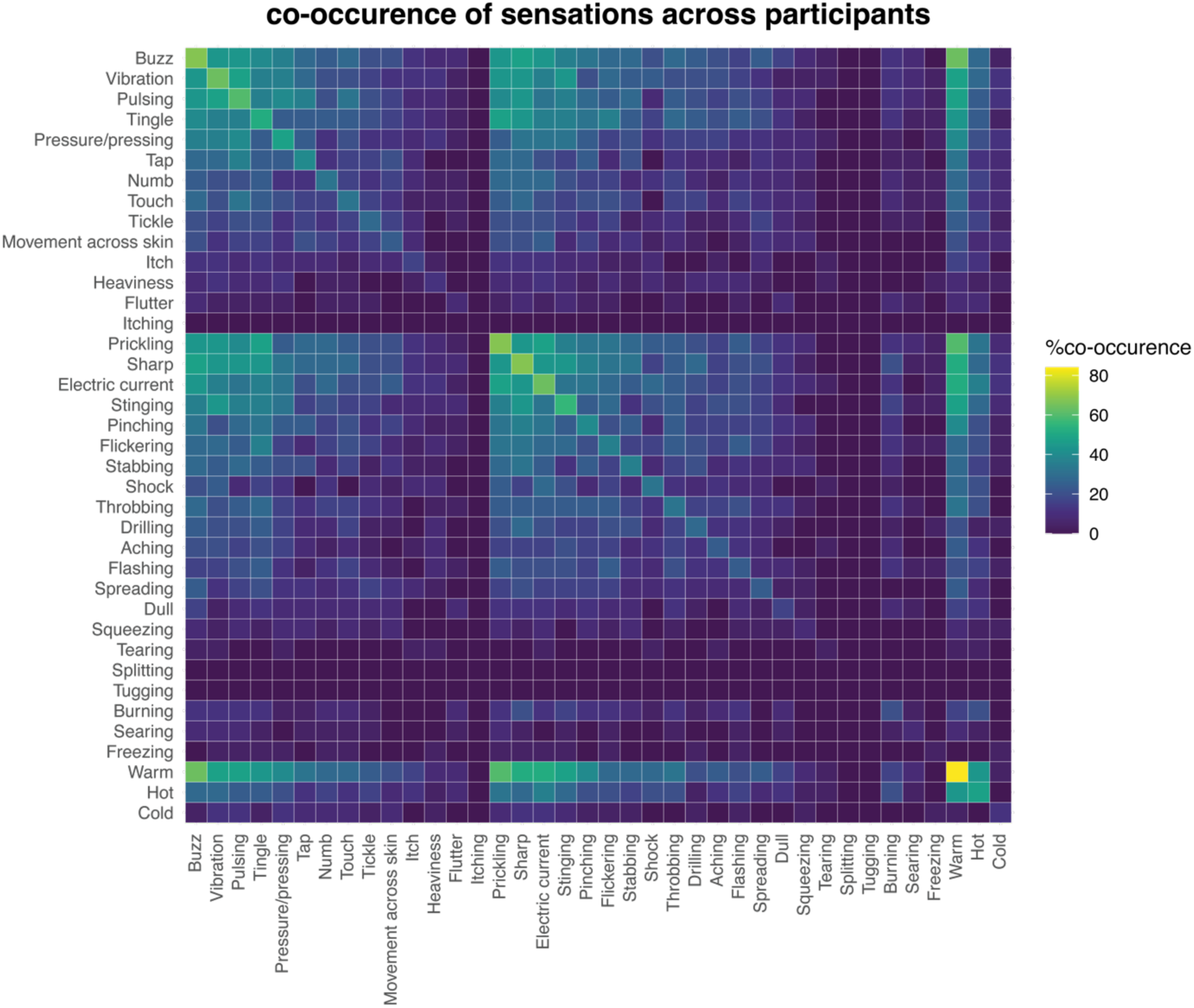
Co-occurrence of somatosensory percepts. Co-occurrence of items on the psychometric questionnaire. Percentages reflect the proportion of participants that felt each pair of sensations. The diagonal depicts the percentage of participants that reported feeling each individual sensation.

**Supplementary Fig. 8.**
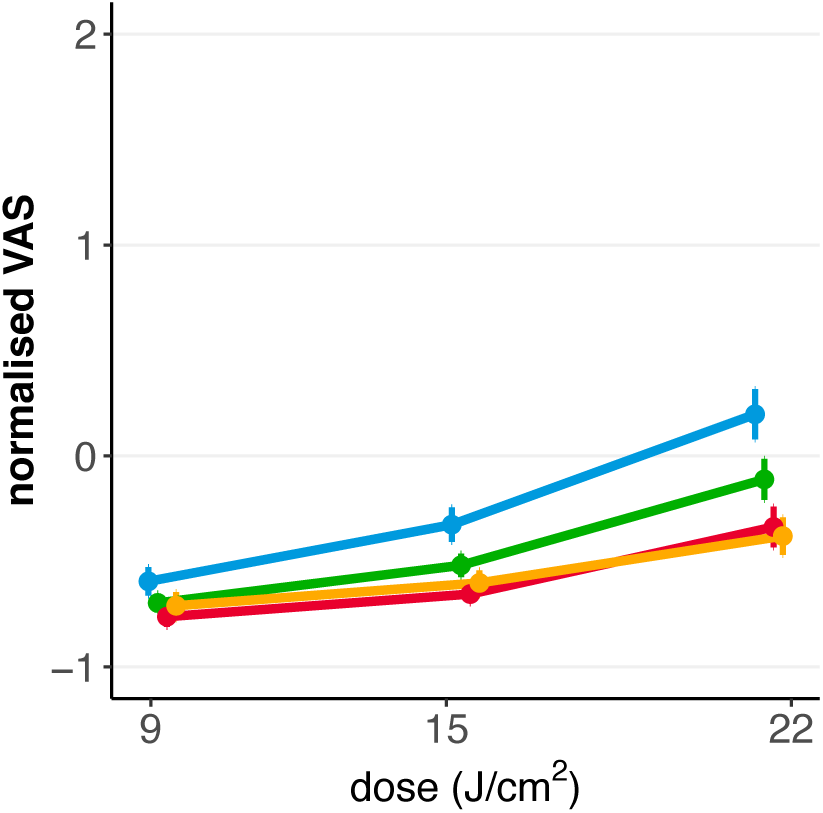
Dose-response (500 kHz) There was a significant e@ect of dose on VAS ratings for 500 kHz TUS. Points represent mean normalised VAS ratings across participants, and error bars depict standard error. Blue = general, green = tactile, orange = thermal, red = painful.

**Supplementary Fig. 9.**
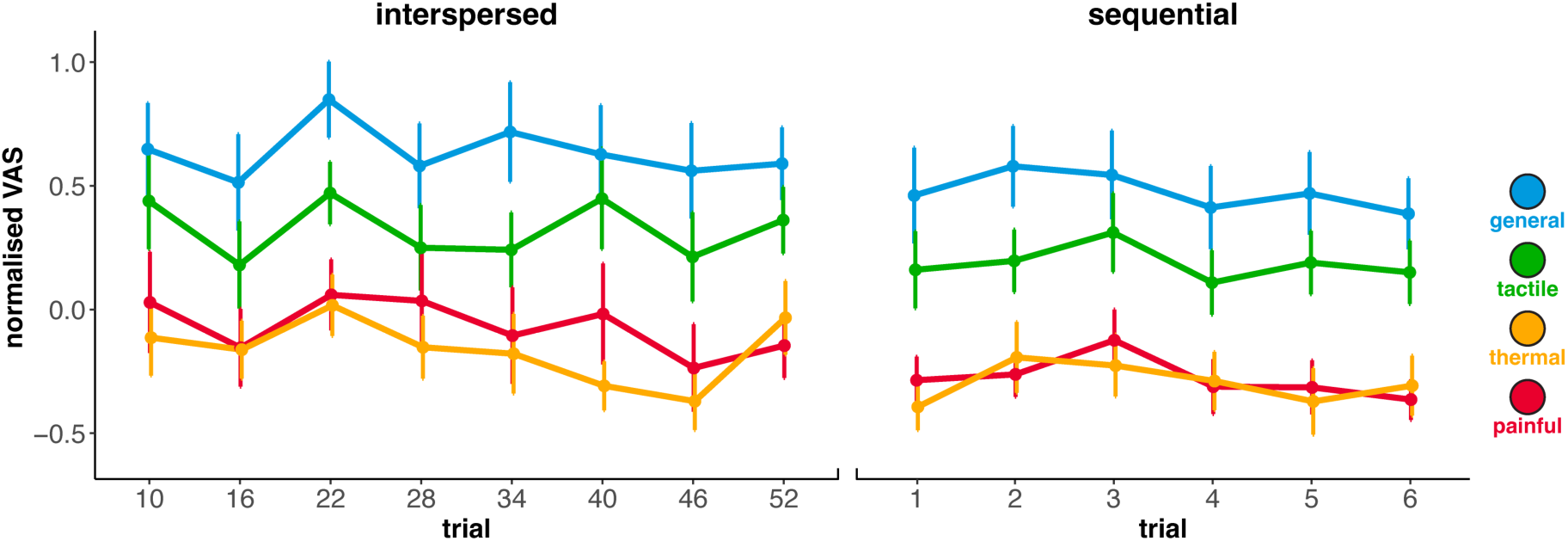
Inter-trial temporal summation. There was no significant e@ect of trial on VAS ratings, with Bayesian analyses providing strong evidence for the null hypothesis (see main text for statistics). This result holds both for identical trials delivered interspersed throughout a block (left) and delivered consecutively (right). Points depict mean normalised VAS ratings; error bars depict standard error.

## References

1. Murphy K, Fouragnan E. The future of transcranial ultrasound as a precision brain interface. PLOS Biology. 2024;22(10):e3002884. doi:10.1371/journal.pbio.3002884

2. Murphy KR, Farrell JS, Bendig J, et al. Optimized ultrasound neuromodulation for non-invasive control of behavior and physiology. Neuron. 2024;112(19):3252–3266.e5. doi:10.1016/j.neuron.2024.07.002

3. Yaakub SN, White TA, Roberts J, et al. Transcranial focused ultrasound-mediated neurochemical and functional connectivity changes in deep cortical regions in humans. Nat Commun. 2023;14(1):5318. doi:10.1038/s41467-023-40998-0

4. Yaakub SN, Bault N, Lojkiewiez M, et al. Non-invasive Ultrasound Deep Neuromodulation of the Human Nucleus Accumbens Increases Win-Stay Behaviour. Published online July 25, 2024:2024.07.25.605068. doi:10.1101/2024.07.25.605068

5. Fouragnan EF, Chau BK, Folloni D, et al. The macaque anterior cingulate cortex translates counterfactual choice value into actual behavioral change. Nature neuroscience. 2019;22(5):797–808.

6. Nakajima K, Osada T, Ogawa A, et al. A causal role of anterior prefrontal-putamen circuit for response inhibition revealed by transcranial ultrasound stimulation in humans. Cell Reports. 2022;40(7):111197. doi:10.1016/j.celrep.2022.111197

7. Bancel T, Béranger B, Daniel M, et al. Sustained reduction of Essential Tremor with low-power non-thermal transcranial focused ultrasound stimulations in humans. Brain Stimulation. Published online May 9, 2024. doi:10.1016/j.brs.2024.05.003

8. Martin E, Roberts M, Grigoras IF, et al. Ultrasound system for precise neuromodulation of human deep brain circuits. Published online June 8, 2024:2024.06.08.597305. doi:10.1101/2024.06.08.597305

9. Riis TS, Losser AJ, Kassavetis P, Moretti P, Kubanek J. Noninvasive modulation of essential tremor with focused ultrasonic waves. J Neural Eng. 2024;21(1):016033. doi:10.1088/1741-2552/ad27ef

10. Webb TD, Wilson MG, Odéen H, Kubanek J. Sustained modulation of primate deep brain circuits with focused ultrasonic waves. Brain Stimulation. 2023;16(3):798–805. doi:10.1016/j.brs.2023.04.012

11. Butler CR, Rhodes E, Blackmore J, et al. Transcranial ultrasound stimulation to human middle temporal complex improves visual motion detection and modulates electrophysiological responses. *Brain Stimulation: Basic*, Translational, and Clinical Research in Neuromodulation. 2022;15(5):1236–1245. doi:10.1016/j.brs.2022.08.022

12. Duecker F, de Graaf TA, Jacobs C, Sack AT. Time- and Task-Dependent Non-Neural Effects of Real and Sham TMS. de Lange FP, ed. PLoS ONE. 2013;8(9):e73813. doi:10.1371/journal.pone.0073813

13. Duecker F, Sack AT. Rethinking the role of sham TMS. Front Psychol. 2015;6. doi:10.3389/fpsyg.2015.00210

14. Kop BR, Shamli Oghli Y, Grippe TC, et al. Auditory confounds can drive online effects of transcranial ultrasonic stimulation in humans. Obleser J, Roiser J, eds. eLife. 2024;12:RP88762. doi:10.7554/eLife.88762

15. Polanía R, Nitsche MA, Ruff CC. Studying and modifying brain function with non-invasive brain stimulation. Nat Neurosci. 2018;21(2):174–187. doi:10.1038/s41593-017-0054-4

16. Fertonani A, Ferrari C, Miniussi C. What do you feel if I apply transcranial electric stimulation? Safety, sensations and secondary induced effects. Clinical Neurophysiology. 2015;126(11):2181–2188. doi:10.1016/j.clinph.2015.03.015

17. Wathra RA, Mulsant BH, Reynolds CF, et al. Differential Placebo Responses for Pharmacotherapy and Neurostimulation in Late-Life Depression. Neuromodulation: Technology at the Neural Interface. 2023;26(8):1585–1591. doi:10.1016/j.neurom.2021.10.019

18. Razza LB, Moffa AH, Moreno ML, et al. A systematic review and meta-analysis on placebo response to repetitive transcranial magnetic stimulation for depression trials. Progress in Neuro-Psychopharmacology and Biological Psychiatry. 2018;81:105–113. doi:10.1016/j.pnpbp.2017.10.016

19. Burke MJ, Kaptchuk TJ, Pascual-Leone A. Challenges of differential placebo effects in contemporary medicine: The example of brain stimulation. Annals of Neurology. 2019;85(1):12–20. doi:10.1002/ana.25387

20. Airan RD, Butts Pauly K. Hearing out Ultrasound Neuromodulation. Neuron. 2018;98(5):875–877. doi:10.1016/j.neuron.2018.05.031

21. Guo H, Hamilton M, Offutt SJ, et al. Ultrasound Produces Extensive Brain Activation via a Cochlear Pathway. Neuron. 2018;98(5):1020–1030.e4. doi:10.1016/j.neuron.2018.04.036

22. Sato T, Shapiro MG, Tsao DY. Ultrasonic Neuromodulation Causes Widespread Cortical Activation via an Indirect Auditory Mechanism. Neuron. 2018;98(5):1031–1041.e5. doi:10.1016/j.neuron.2018.05.009

23. Braun V, Blackmore J, Cleveland RO, Butler CR. Transcranial ultrasound stimulation in humans is associated with an auditory confound that can be effectively masked. Brain Stimulation. 2020;13(6):1527–1534. doi:10.1016/j.brs.2020.08.014

24. Johnstone A, Nandi T, Martin E, Bestmann S, Stagg C, Treeby B. A range of pulses commonly used for human transcranial ultrasound stimulation are clearly audible. Brain Stimulation. 2021;14(5):1353–1355. doi:10.1016/j.brs.2021.08.015

25. Liang W, Guo H, Mittelstein DR, Shapiro MG, Shimojo S, Shehata M. Auditory Mondrian masks the airborne-auditory artifact of focused ultrasound stimulation in humans. *Brain Stimulation: Basic*, Translational, and Clinical Research in Neuromodulation. 2023;16(2):604–606. doi:10.1016/j.brs.2023.03.002

26. Choi MH, Li N, Popelka G, Butts Pauly K. Development and validation of a computational method to predict unintended auditory brainstem response during transcranial ultrasound neuromodulation in mice. Brain Stimulation. 2023;16(5):1362–1370. doi:10.1016/j.brs.2023.09.004

27. Mohammadjavadi M, Ye PP, Xia A, Brown J, Popelka G, Pauly KB. Elimination of peripheral auditory pathway activation does not affect motor responses from ultrasound neuromodulation. Brain Stimul. 2019;12(4):901–910. doi:10.1016/j.brs.2019.03.005

28. Brinker ST, Preiswerk F, White PJ, Mariano TY, McDannold NJ, Bubrick EJ. Focused Ultrasound Platform for Investigating Therapeutic Neuromodulation Across the Human Hippocampus. Ultrasound in Medicine & Biology. 2020;46(5):1270–1274. doi:10.1016/j.ultrasmedbio.2020.01.007

29. Spivak NM, Sanguinetti JL, Monti MM. Focusing in on the Future of Focused Ultrasound as a Translational Tool. Brain Sciences. 2022;12(2):158. doi:10.3390/brainsci12020158

30. Murphy KR, Good C, Di Ianni T, et al. Head-Wearable Devices for Positioning Ultrasound Transducers for Brain Stimulation. Published online June 1, 2023. Accessed December 4, 2024. https://patentscope.wipo.int/search/en/detail.jsf?docId=WO2023097122

31. Smith CS, O’Driscoll C, Ebbini ES. Spatio-Spectral Ultrasound Characterization of Reflection and Transmission Through Bone With Temperature Dependence. *IEEE Transactions on Ultrasonics*, Ferroelectrics, and Frequency Control. 2022;69(5):1727–1737. doi:10.1109/TUFFC.2022.3163225

32. Hoang-Dang B, Halavi SE, Rotstein NM, et al. Transcranial Focused Ultrasound Targeting the Amygdala May Increase Psychophysiological and Subjective Negative Emotional Reactivity in Healthy Older Adults. Biological Psychiatry Global Open Science. 2024;4(5):100342. doi:10.1016/j.bpsgos.2024.100342

33. Fan JM, Woodworth K, Murphy KR, et al. Thalamic transcranial ultrasound stimulation in treatment resistant depression. Brain Stimulation. 2024;17(5):1001–1004. doi:10.1016/j.brs.2024.08.006

34. L. Gavrilov. Focused ultrasound stimulation of the peripheral nervous system: physical basis and practical applications. Int J Mod Phys Adv Theory Appl. 2016;1(1):45–118.

35. Riis T, Kubanek J. Effective Ultrasonic Stimulation in Human Peripheral Nervous System. IEEE Transactions on Biomedical Engineering. 2022;69(1):15–22. doi:10.1109/TBME.2021.3085170

36. Qin L, Dou M, Niu L, et al. Low-Intensity Focused Ultrasound Stimulation on Fingertip Can Evoke Fine Tactile Sensations and Different Local Hemodynamic Responses. IEEE Transactions on Neural Systems and Rehabilitation Engineering. 2024;32:4086–4097. doi:10.1109/TNSRE.2024.3493925

37. Muratore R, Vaitekunas J. Ultrasonic Bioeffects on Peripheral Nerves. Acoustics Today. 2012;8:38. doi:10.1121/1.4788650

38. Dalecki D, Child SZ, Raeman CH, Carstensen EL. Tactile perception of ultrasound. The Journal of the Acoustical Society of America. 1995;97(5):3165–3170. doi:10.1121/1.411877

39. Dubin AE, Patapoutian A. Nociceptors: the sensors of the pain pathway. J Clin Invest. 2010;120(11):3760–3772. doi:10.1172/JCI42843

40. Griensven H van, Strong J, Unruh AM. Pain E-Book: A Textbook for Therapists. Elsevier Health Sciences; 2013.

41. Gavrilov LR, Tsirulnikov EM, Davies I ab I. Application of focused ultrasound for the stimulation of neural structures. Ultrasound in Medicine & Biology. 1996;22(2):179–192. doi:10.1016/0301-5629(96)83782-3

42. Lee W, Kim H, Lee S, Yoo SS, Chung YA. Creation of various skin sensations using pulsed focused ultrasound: Evidence for functional neuromodulation. International Journal of Imaging Systems and Technology. 2014;24(2):167–174. doi:10.1002/ima.22091

43. Darmani G, Bergmann TO, Butts Pauly K, et al. Non-invasive transcranial ultrasound stimulation for neuromodulation. Clinical Neurophysiology. 2022;135:51–73. doi:10.1016/j.clinph.2021.12.010

44. Sorum B, Rietmeijer RA, Gopakumar K, Adesnik H, Brohawn SG. Ultrasound activates mechanosensitive TRAAK K+ channels through the lipid membrane. Proc Natl Acad Sci U S A. 2021;118(6):e2006980118. doi:10.1073/pnas.2006980118

45. Prieto ML, Firouzi K, Khuri-Yakub BT, Maduke M. Activation of Piezo1 but Not NaV1.2 Channels by Ultrasound at 43 MHz. Ultrasound Med Biol. 2018;44(6):1217–1232. doi:10.1016/j.ultrasmedbio.2017.12.020

46. Kubanek J, Shi J, Marsh J, Chen D, Deng C, Cui J. Ultrasound modulates ion channel currents. Sci Rep. 2016;6(1):24170. doi:10.1038/srep24170

47. Zhu J, Xian Q, Hou X, et al. The mechanosensitive ion channel Piezo1 contributes to ultrasound neuromodulation. Proceedings of the National Academy of Sciences. 2023;120(18):e2300291120. doi:10.1073/pnas.2300291120

48. Jerusalem A, Al-Rekabi Z, Chen H, et al. Electrophysiological-mechanical coupling in the neuronal membrane and its role in ultrasound neuromodulation and general anaesthesia. Acta Biomaterialia. 2019;97:116–140. doi:10.1016/j.actbio.2019.07.041

49. Xu T, Zhang Y, Li D, Lai C, Wang S, Zhang S. Mechanosensitive Ion Channels Piezo1 and Piezo2 Mediate Motor Responses In Vivo During Transcranial Focused Ultrasound Stimulation of the Rodent Cerebral Motor Cortex. IEEE Transactions on Biomedical Engineering. 2024;71(10):2900–2910. doi:10.1109/TBME.2024.3401136

50. Xu K, Yang Y, Hu Z, et al. TRPV1-mediated sonogenetic neuromodulation of motor cortex in freely moving mice. J Neural Eng. 2023;20(1):016055. doi:10.1088/1741-2552/acbba0

51. Davies I ab I, Gavrilov LR, Tsirulnikov EM. Application of focused ultrasound for research on pain. PAIN. 1996;67(1):17–27. doi:10.1016/0304-3959(96)03042-4

52. Gavrilov LR. Use of focused ultrasound for stimulation of nerve structures. Ultrasonics. 1984;22(3):132–138. doi:10.1016/0041-624x(84)90008-8

53. Gavrilov LR, Tsirulnikov EM. Focused ultrasound as a tool to input sensory information to humans (Review). Acoust Phys. 2012;58(1):1–21. doi:10.1134/S1063771012010083

54. Kop B, Verhagen L, Ouden H den. Somatosensory Confounds of Transcranial Ultrasound Stimulation. Published online September 6, 2024. doi:10.17605/OSF.IO/UMTZD

55. Treeby BE, Cox BT. k-Wave: MATLAB toolbox for the simulation and reconstruction of photoacoustic wave fields. J Biomed Opt. 2010;15(2):021314. doi:10.1117/1.3360308

56. Miscouridou M, Pineda-Pardo JA, Stagg CJ, Treeby BE, Stanziola A. Classical and Learned MR to Pseudo-CT Mappings for Accurate Transcranial Ultrasound Simulation. IEEE Trans Ultrason Ferroelectr Freq Control. 2022;69(10):2896–2905. doi:10.1109/TUFFC.2022.3198522

57. Martin E, Aubry JF, Schafer M, Verhagen L, Treeby B, Pauly KB. ITRUSST Consensus on Standardised Reporting for Transcranial Ultrasound Stimulation. *ArXiv*. Published online February 15, 2024:arXiv:2402.10027v1.

58. Klein-fiügge MC, Fouragnan EF, Martin E. The importance of acoustic output measurement and monitoring for the replicability of transcranial ultrasonic stimulation studies. *Brain Stimulation: Basic*, Translational, and Clinical Research in Neuromodulation. 2024;17(1):32–34. doi:10.1016/j.brs.2023.12.002

59 . Murphy KR, Nandi T, Kop B, et al. A Practical Guide to Transcranial Ultrasonic Stimulation from the IFCN-endorsed ITRUSST Consortium. Published online September 20, 2024. doi:10.48550/arXiv.2407.07646

60. Holmes NP, ed. Somatosensory Research Methods. Humana Press; 2023.

61. Taylor MM, Creelman CD. PEST: Efficient Estimates on Probability Functions. The Journal of the Acoustical Society of America. 1967;41(4A):782–787. doi:10.1121/1.1910407

62. Kim LH, McLeod RS, Kiss ZHT. A new psychometric questionnaire for reporting of somatosensory percepts. J Neural Eng. 2018;15(1):013002. doi:10.1088/1741-2552/aa966a

63. Peirce J, Gray JR, Simpson S, et al. PsychoPy2: Experiments in behavior made easy. Behav Res. 2019;51(1):195–203. doi:10.3758/s13428-018-01193-y

64. Aubry JF, Attali D, Schafer M, et al. ITRUSST Consensus on Biophysical Safety for Transcranial Ultrasonic Stimulation. Published online July 12, 2024. doi:10.48550/arXiv.2311.05359

65. Nandi T, Kop BR, Naftchi-Ardebili K, Stagg CJ, Pauly KB, Verhagen L. Biophysical effects and neuromodulatory dose of transcranial ultrasonic stimulation. Brain Stimulation. Published online March 5, 2025. doi:10.1016/j.brs.2025.02.019

66. Fong PY, Kop BR, Evans C, et al. Failed Double-Blind Replication of Offline 5Hz-rTUS-Induced Corticospinal Excitability. Published online November 26, 2024:2024.11.25.625187. doi:10.1101/2024.11.25.625187

67. Barr DJ, Levy R, Scheepers C, Tily HJ. Random effects structure for confirmatory hypothesis testing: Keep it maximal. Journal of Memory and Language. 2013;68(3):255–278. doi:10.1016/j.jml.2012.11.001

68. Bates D, Mächler M, Bolker B, Walker S. Fitting Linear Mixed-Effects Models Using lme4. J Stat Soft. 2015;67(1). doi:10.18637/jss.v067.i01

69. Djouhri L, Lawson SN. Abeta-fiber nociceptive primary afferent neurons: a review of incidence and properties in relation to other afferent A-fiber neurons in mammals. Brain Res Brain Res Rev. 2004;46(2):131–145. doi:10.1016/j.brainresrev.2004.07.015

70. Nandi T, Kop BR, Naftchi-Ardebili K, Stagg CJ, Pauly KB, Verhagen L. Biophysical effects and neuromodulatory dose of transcranial ultrasonic stimulation. *ArXiv*. Published online July 3, 2024:arXiv:2406.19869v3.

71. Nandi T, Kop BR, Butts Pauly K, Stagg CJ, Verhagen L. The relationship between parameters and effects in transcranial ultrasonic stimulation. Brain Stimulation. 2024;17(6):1216–1228. doi:10.1016/j.brs.2024.10.008

72. Kubanek J, Shukla P, Das A, Baccus SA, Goodman MB. Ultrasound Elicits Behavioral Responses through Mechanical Effects on Neurons and Ion Channels in a Simple Nervous System. J Neurosci. 2018;38(12):3081–3091. doi:10.1523/JNEUROSCI.1458-17.2018

73. Yoo S, Mittelstein DR, Hurt RC, Lacroix J, Shapiro MG. Focused ultrasound excites cortical neurons via mechanosensitive calcium accumulation and ion channel amplification. Nat Commun. 2022;13(1):493. doi:10.1038/s41467-022-28040-1

74. Nandi T, Johnstone A, Martin E, et al. Ramped V1 transcranial ultrasonic stimulation modulates but does not evoke visual evoked potentials. *Brain Stimulation: Basic*, Translational, and Clinical Research in Neuromodulation. 2023;16(2):553–555. doi:10.1016/j.brs.2023.02.004

75. Zadeh AK, Raghuram H, Shrestha S, et al. The effect of transcranial ultrasound pulse repetition frequency on sustained inhibition in the human primary motor cortex: A double-blind, sham-controlled study. Brain Stimulation. 2024;17(2):476–484. doi:10.1016/j.brs.2024.04.005

76. Deffieux T, Younan Y, Wattiez N, Tanter M, Pouget P, Aubry JF. Low-Intensity Focused Ultrasound Modulates Monkey Visuomotor Behavior. Current Biology. 2013;23(23):2430–2433. doi:10.1016/j.cub.2013.10.029

77. Pauly KB, Kop B, Qui Z, et al. Transcranial ultrasound stimulation: considerations for pulse shaping. *Brain Stimulation: Basic*, Translational, and Clinical Research in Neuromodulation. 2023;16(1):200–201. doi:10.1016/j.brs.2023.01.258

78. Yu K, Niu X, Krook-Magnuson E, He B. Intrinsic functional neuron-type selectivity of transcranial focused ultrasound neuromodulation. Nature Communications. 2021;12. doi:10.1038/s41467-021-22743-7

79. Lundström R, Strömberg T, Lundborg G. Vibrotactile perception threshold measurements for diagnosis of sensory neuropathy. Int Arch Occup Environ Heath. 1992;64(3):201–207. doi:10.1007/BF00380910

80. Mahns DA, Perkins NM, Sahai V, Robinson L, Rowe MJ. Vibrotactile Frequency Discrimination in Human Hairy Skin. Journal of Neurophysiology. 2006;95(3):1442–1450. doi:10.1152/jn.00483.2005

81. Kubanek J, Brown J, Ye P, Pauly KB, Moore T, Newsome W. Remote, brain region–specific control of choice behavior with ultrasonic waves. Sci Adv. 2020;6(21):eaaz4193. doi:10.1126/sciadv.aaz4193

82. Samuel N, Ding MYR, Sarica C, et al. Accelerated Transcranial Ultrasound Neuromodulation in Parkinson’s Disease: A Pilot Study. Movement Disorders. 2023;38(12):2209–2216. doi:10.1002/mds.29622

83. Riis TS, Feldman DA, Losser AJ, Okifuji A, Kubanek J. Noninvasive targeted modulation of pain circuits with focused ultrasonic waves. PAIN. 2024;165(12):2829. doi:10.1097/j.pain.0000000000003322

84. Riis TS, Feldman DA, Kwon SS, et al. Noninvasive modulation of subcallosal cingulate and depression with focused ultrasonic waves. Biological Psychiatry. Published online October 2024:S0006322324016627. doi:10.1016/j.biopsych.2024.09.029

85. Lee SA, Kamimura HAS, Burgess MT, Konofagou EE. Displacement Imaging for Focused Ultrasound Peripheral Nerve Neuromodulation. IEEE Transactions on Medical Imaging. 2020;39(11):3391–3402. doi:10.1109/TMI.2020.2992498

86. Menz MD, Ye P, Firouzi K, et al. Radiation Force as a Physical Mechanism for Ultrasonic Neurostimulation of the *Ex Vivo* Retina. J Neurosci. 2019;39(32):6251–6264. doi:10.1523/JNEUROSCI.2394-18.2019

87. Kim YH, Kang KC, Kim JN, et al. Patterned Interference Radiation Force for Transcranial Neuromodulation. Ultrasound in Medicine & Biology. 2022;48(3):497–511. doi:10.1016/j.ultrasmedbio.2021.11.006

88. Kerstens S, Orban de Xivry JJ, Mc Laughlin M. A novel tDCS control condition using optimized anesthetic gel to block peripheral nerve input. Front Neurol. 2022;13. doi:10.3389/fneur.2022.1049409

## References

1. Martin E, Aubry JF, Schafer M, Verhagen L, Treeby B, Pauly KB. ITRUSST Consensus on Standardised Reporting for Transcranial Ultrasound Stimulation. *ArXiv*. Published online February 15, 2024:arXiv:2402.10027v1.

2. Aubry JF, Attali D, Schafer M, et al. ITRUSST Consensus on Biophysical Safety for Transcranial Ultrasonic Stimulation. Published online July 12, 2024. doi:10.48550/arXiv.2311.05359

3. Haeussinger FB, Heinzel S, Hahn T, Schecklmann M, Ehlis AC, Fallgatter AJ. Simulation of Near-Infrared Light Absorption Considering Individual Head and Prefrontal Cortex Anatomy: Implications for Optical Neuroimaging. PLOS ONE. 2011;6(10):e26377. doi:10.1371/journal.pone.0026377

4. Light AE. Histological study of human scalps exhibiting various degrees of non-specific baldness. J Invest Dermatol. 1949;13(2):53–59. doi:10.1038/jid.1949.67

5. Garn SM, Selby S, Young R. SCALP THICKNESS AND THE FAT-LOSS THEORY OF BALDING. AMA Archives of Dermatology and Syphilology. 1954;70(5):601–608. doi:10.1001/archderm.1954.01540230051006

6. Treeby BE, Cox BT. k-Wave: MATLAB toolbox for the simulation and reconstruction of photoacoustic wave fields. J Biomed Opt. 2010;15(2):021314. doi:10.1117/1.3360308

7. Miscouridou M, Pineda-Pardo JA, Stagg CJ, Treeby BE, Stanziola A. Classical and Learned MR to Pseudo-CT Mappings for Accurate Transcranial Ultrasound Simulation. IEEE Trans Ultrason Ferroelectr Freq Control. 2022;69(10):2896–2905. doi:10.1109/TUFFC.2022.3198522

8. Leek MR. Adaptive procedures in psychophysical research. Perception & Psychophysics. 2001;63(8):1279–1292. doi:10.3758/BF03194543

9. Holmes NP, ed. Somatosensory Research Methods. Humana Press; 2023.

10. Taylor MM, Creelman CD. PEST: Efficient Estimates on Probability Functions. The Journal of the Acoustical Society of America. 1967;41(4A):782–787. doi:10.1121/1.1910407

